# Snail induces epithelial cell extrusion through transcriptional control of RhoA contractile signaling and cell matrix adhesion

**DOI:** 10.1101/621698

**Authors:** Kenneth Wee, Suroor Zadeh, Suzie Verma, Roger J. Daly, Alpha S. Yap, Melissa J. Davis, Srikanth Budnar

**Affiliations:** Division of Cell Biology and Molecular Medicine, Institute for Molecular Bioscience, The University of Queensland, St. Lucia, Brisbane, Queensland, Australia 4072; Bioinformatics Division, Walter and Eliza Hall Institute of Medical Research, 1G Royal Parade, Parkville, VIC 3052, Australia; Cancer Program, Biomedicine Discovery Institute and Department of Biochemistry and Molecular Biology, Monash University

**Keywords:** Snail, RhoA, contractility, extrusion

## Abstract

Cell extrusion is a morphogenetic process that is implicated in epithelial homeostasis and highly dependent on the cellular insult and context. Minorities of cells expressing HRas^V12^ or undergoing apoptosis are typically extruded apically in vertebrate cells. However, basal extrusion (delamination) predominates when mammalian cells express oncogenic KRas and during development in Drosophila. To explore if the morphogenetic transcription factor, Snail, induces extrusion, we inducibly expressed a metabolically stabilized Snail^6SA^ transgene mosaically in confluent MCF-7 monolayers. We found that the morphogenetic impact of Snail^6SA^ was critically influenced by the proportion of cells in which it was expressed. When expressed in small clusters (<3 cells) within confluent monolayers, Snail^6SA^ expression induced apical cell extrusion. In contrast, confluent cultures of Snail^6SA^ expressing cells were retained in the monolayer to eventually undergo basal extrusion (delamination). Transcriptomic profiling revealed that Snail^6SA^ did not substantively alter the balance of epithelial:mesenchymal genes. However, we identified a gene transcriptional network that led to the upregulation of RhoA signalling, actomyosin contractility and reduced basal adhesion in Snail^6SA^ expressing cells. We show that this was necessary to drive both apical extrusion and basal delamination. Thus, contractile upregulation by RhoA along with weakened basal adhesion provides a pathway for Snail to influence epithelial morphogenesis independently of classic Epithelial to Mesenchymal Transition (EMT).

## Introduction

Transcription factors of the Snail family (Snail TFs) are important regulators of epithelial morphogenesis both during development and in post-developmental life (Nieto 2002). Its morphogenetic impact has commonly been ascribed to the ability of Snail to promote epithelial-to-mesenchymal transitions (EMT) (Peinado, Olmeda et al. 2007, Lamouille, Xu et al. 2014, Nieto, Huang et al. 2016, Dongre and Weinberg 2018). For example, during gastrulation in sea urchin embryos, Snail is essential for the EMT of epithelial cells into primary mesenchymal cells leading to their delamination and invasion into the blastocoele (Wu and McClay 2007). Similarly, Snail has been implicated in tumor invasiveness. EMT typically involves the repression of genes associated with the epithelial phenotype, such as E-cadherin, and the expression of mesenchymal genes, such as vimentin and N-cadherin (Batlle, Sancho et al. 2000, Cano, Perez-Moreno et al. 2000). In this model, downregulation of E-cadherin is proposed to allow dissociation of cells through loss of cell-cell junctions that facilitates their invasion into stroma, thereby reflecting the morphogenetic impact of Snail during cellular transdifferentiation.

Snail is also implicated in morphogenetic events where cellular contractility is a major driving process. During *Drosophila* gastrulation, formation of the ventral furrow is mediated by contractility in the cortical actomyosin cytoskeleton that is regulated by both Snail and another transcription factor, Twist (Martin, Kaschube et al. 2009, Weng and Wieschaus 2016). Twist causes localized activation of Rho1 through the concerted action of its targets, Fog and T48 (Costa, Wilson et al. 1994, Nikolaidou and Barrett 2004, Kolsch, Seher et al. 2007). Twist also induces the expression of Snail in these ventral cells, which is responsible for the pulsed contractions of medial-apical acto-myosin networks that constrict the apical poles of cells and drive formation of the ventral furrow (Martin, Kaschube et al. 2009). Actomyosin-based cortical contractility has also been implicated in basal delamination of cells, both during early mouse development and in *C. elegans* gastrulation (Pohl, Tiongson et al. 2012). This is coordinated with cell-to-cell adherens junctions (Roh-Johnson, Shemer et al. 2012). Indeed, both cortical contractility and E-cadherin were necessary for Snail-induced apical constriction during *Drosophila* gastrulation (Weng and Wieschaus 2016). Thus, regulation of cellular contractility provides an alternative pathway for Snail to influence epithelial organization.

In this report, we analyse the ability of Snail to promote the morphogenetic phenomenon of epithelial cell extrusion. This is a ubiquitous event, where minorities of cells are expelled from the epithelium either apically, towards the lumen (apical extrusion), or in a basal direction into the stroma (basal extrusion, also known as delamination). Extrusion is a biomechanical phenomenon that can be induced by diverse cellular changes, ranging from apoptosis to tissue over-crowding. However, our understanding of the mechanisms that drive extrusion remains incomplete. Nor do we understand what determines the direction of extrusion, although a role for cortical capture of microtubules has been identified (Slattum, Mcgee et al. 2009). Here we show that expression of a stabilized Snail transgene (Snail^6SA^) activates cell contractility through a transcriptional network that stimulates the RhoA GTPase. This hyper-contractile cortex induces apical extrusion upon mosaic expression, when Snail^6SA^ cells are surrounded by wild-type (WT) epithelia. In contrast, when expressed in large groups of cells, Snail^6SA^ induces delamination due to transcriptional suppression of integrin-based adhesion coupled to a hyper-contractile cortex. This represents a hitherto unrecognized phenomenon of epithelial behaviour driven by Snail, independent of its canonical role in EMT.

## Results

### Cell-proportions determine the direction of extrusion of Snail expressing cells

We developed an MCF-7 cell line that inducibly expresses Snail^6SA^, a stabilized mutant that is less susceptible to metabolic turnover (Zhou, Deng et al. 2004). Characteristically, Snail^6SA^ expression was evident within 6 hrs of doxycycline induction which increased to ∼3 fold at 48 hrs (Figure S1A and B.) Strikingly, we found that when expressed in small clusters (< 3 cells) within otherwise confluent WT monolayers, Snail^6SA^ cells underwent apical extrusion from the epithelium within 48 hr of induction (Figure 1A and B; Figure S1C and D). Such apical extrusion has also been described when epithelial cells undergo apoptosis (Michael, Meiring et al. 2016). However, extruded Snail^6SA^ cells showed no evidence of apoptosis, as identified by staining for annexin V (Figure S1E and F). Similar apical extrusion was observed when Snail^6SA^ was expressed in MCF-7 cells that had been CRISPR/Cas9-engineered to express E-cadherin-GFP (MCF-7^GFP-E-cad^) (Figure 1C, S1K - Q and Supplemental Movie 1) and in Caco-2 cells (Figure S1G - J). Thus, apical extrusion appeared to be a common epithelial response to the mosaic expression of Snail^6SA^.

**Figure 1.**
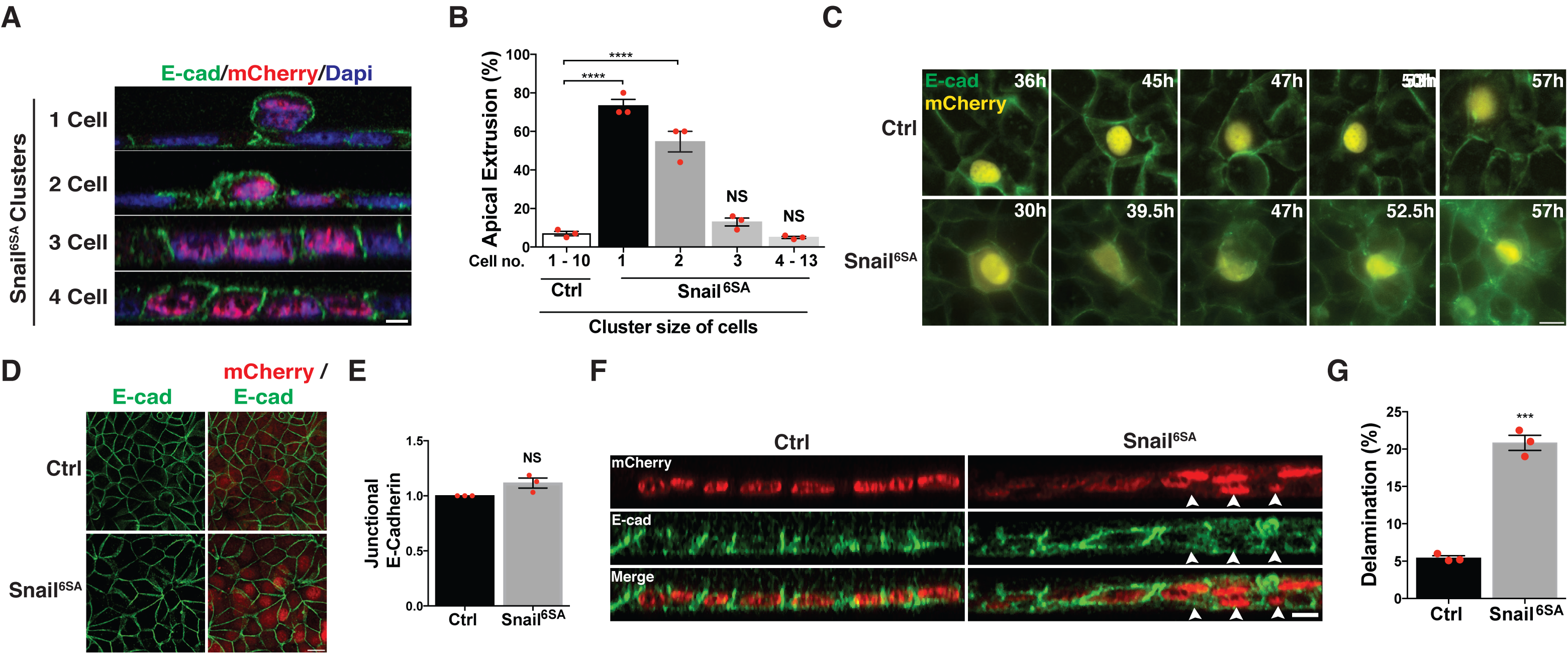
Cell-proportions determine the direction of extrusion of Snail^6SA^ expressing cells. **A, B)** Representative XZ images **(A)** and quantification **(B)** of apically extruded Snail^6SA^ cell clusters when mixed with wild type parental MCF-7 cells, immunostained with E-cadherin and mCherry. **C)** Time-lapse images single cells expressing either mCherry^NLS^ (Ctrl) or Snail^6SA^ within a confluent monolayer of MCF-7 cells in which the endogenous E-cadherin is tagged with GFP. Time in hours represents induction of mCherry^NLS^ (Ctrl) or Snail^6SA^ expression with doxycycline. **D, E)** Representative images **(D)** and quantification **(E)** of MCF-7 cells stably expressing either mCherry^NLS^ (Ctrl) or Snail^6SA^ immunostained with E-cadherin and mCherry. **F, G)** Representative XZ images **(F)** and quantification **(G)** of delaminated mCherry^NLS^ (Ctrl) or Snail^6SA^ expressing cells, immunostained with E-cadherin and mCherry. Arrows indicate delaminated cells. All mCherry^NLS^ (Ctrl) and Snail^6SA^ cells were treated with 3μg/mL of doxycycline for 48 hours. Data represent means ± S.E.M (*p < 0.05, **p < 0.01, ***p < 0.001 and ns = not significant) and n = 3 independent experiments; Students t-test (B, E, G). Scale bars in XY and XZ images represent 10 μm and 5 μm respectively.

However, the morphogenetic impact of Snail^6SA^ depended on the number of cells that expressed the transgene within the epithelium. We found that when Snail^6SA^ was expressed in clusters > 3 cells within the monolayer, they were retained in the epithelium and did not undergo apical extrusion (Figure 1A and B). Indeed, monolayers that uniformly expressed Snail^6SA^ retained E-cadherin-based cell-cell junctions and overall epithelial integrity (Figure 1D and E). However, closer inspection revealed the presence of Snail^6SA^ cells underneath the primary layer of the epithelium (Figure 1F and G). Furthermore, live imaging of Snail^6SA^ monolayers showed cells losing their apical surface to basally exit the monolayer (Supplemental Movie 2). This suggested that Snail^6SA^ expressing cells could ultimately undergo basal delamination (also described as basal extrusion) if they escape apical extrusion.

### Snail^6SA^ cells maintain epithelial identity

We next sought out to understand the changes in signalling downstream of Snail^6SA^ that underpin these cell behaviours. For this, we compared the transcriptional profiles of control monolayers and Snail^6A^ monolayers. RNA-seq analysis performed on total RNA extracted from 3 independent monolayer cultures identified 8843 differentially expressed genes (4280 Down-regulated, 4563 Up-regulated, FDR < 0.05). In addition to the expected increase in Snail (P = 1.06 × 10^-21^, FDR = 5.882478e-18, logFC = 6.04), the EMT-inducing transcription factor ZEB1 was also upregulated (P = 5.9 × 10^-9^, FDR = 8.57E-08, logFC=2.23) (Figure S2A). The differentially expressed genes were examined for changes in epithelial- and mesenchymal-specific genes using the framework of Tan et al. (Tan, Miow et al. 2014). Surprisingly, we found no significant differences in the expression of genes associated with EMT, including the canonical markers E-cadherin and Vimentin. Western analysis confirmed that E-cadherin and vimentin protein expression was not altered in Snail^6A^ cells, compared with control monolayers (Figure 2C). Overall, this suggested that Snail^6SA^ did not induce EMT in these cells.

**Figure 2.**
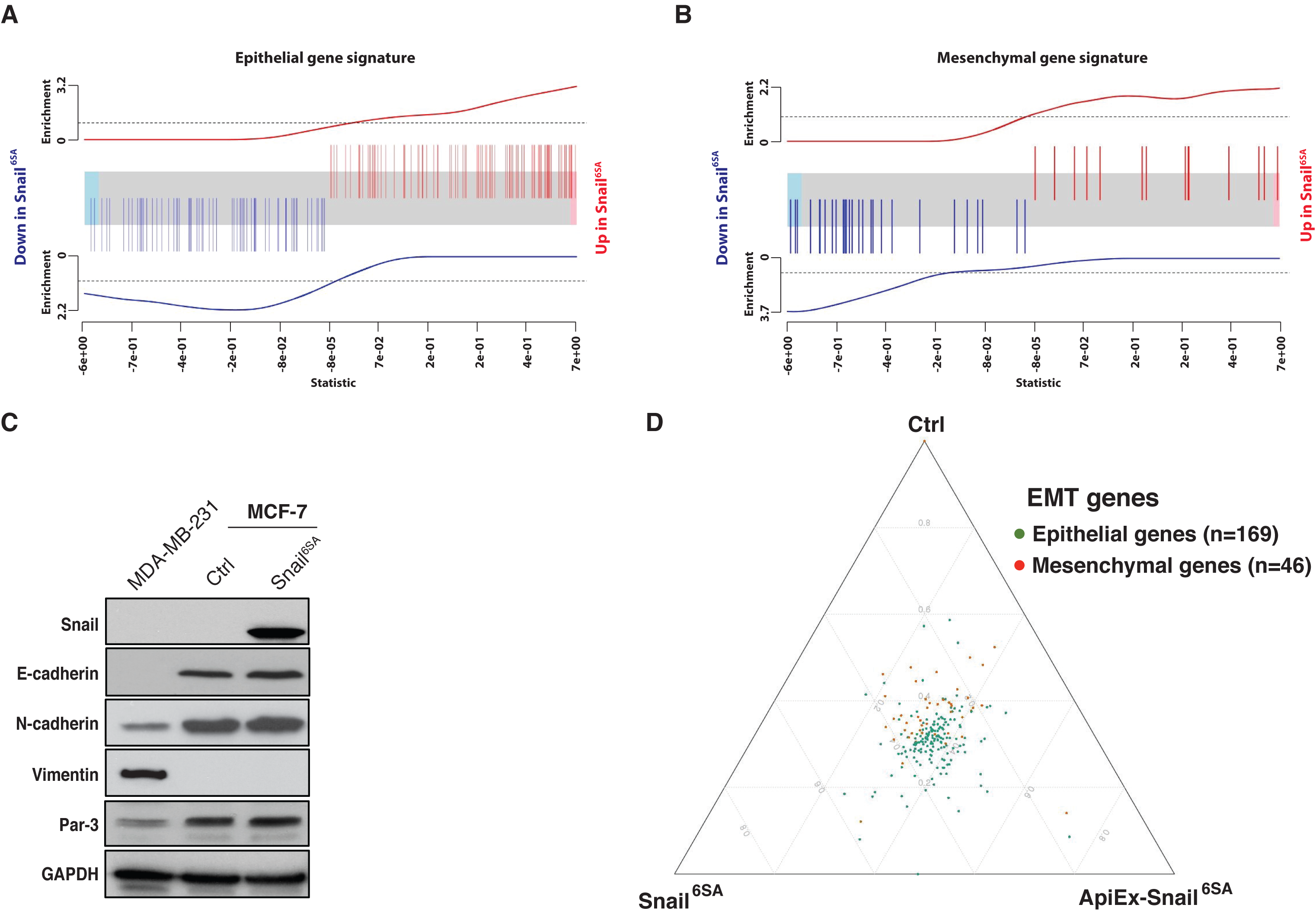
Snail^6SA^ cells maintain epithelial identity. (**A, B**) Enrichment plots for epithelial (**A**) and mesenchymal (**B**) gene sets. **A**) The epithelial signature genes remain strongly expressed and upregulated in Snail^65A^ samples, with a strong concentration of upregulated epithelial genes (strong upward trend in the red line). **B**) A small number of mesenchymal genes are also upregulated, however there is a stronger concentration of downregulated mesenchymal genes (strong downwards trend in the blue line), indicating no strong shift to a mesenchymal molecular phenotype. **C)** Immunoblots of total Snail, E-cadherin, N-cadherin, Vimentin, Par-3 and GAPDH in MDA- MB-231 and MCF-7 cells stably expressing mCherry^NLS^ (Ctrl) or Snail^6SA^. **D)** Ternary plot representing the relative abundance of epithelial (green) and mesenchymal (red) genes between mCherry^NLS^ (Ctrl), Snail^6SA^ and apically extruded Snail^6SA^ (ApiEx- Snail^6SA^) cells. Genes localizing in the centre of the plot have similar relative abundances across three conditions, while genes that shift towards a specific condition indicate relatively higher abundance. All mCherry^NLS^ (Ctrl) and Snail^6SA^ cells were treated with 3μg/mL of doxycycline for 48 hours.

To corroborate these findings, we used the epithelial and mesenchymal signature reported by Tan et al., to score the individual RNA-Seq samples for control and Snail^6SA^ monolayers as well as for apically-extruded Snail^6SA^ cells (ApiEx-Snail^6SA^) (Tan, Miow et al. 2014). The epithelial and mesenchymal scores for these samples were plotted against the Epithelial/Mesenchymal scores in TCGA, along with those for CCLE breast cancer cell lines (Figure 2D). The comparison of these scores revealed that all the samples for Snail^6SA^ and ApiEx-Snail^6SA^ showed a strong tendency to maintain epithelial identity. Together, these results indicate that Snail^6SA^ cells maintain epithelial identity whether in monolayers or even after being apically extruded. This suggested that apical extrusion of Snail^6SA^ cells was not attributable to their having undergone EMT.

### RhoA contractile signalling pathway is hyper-activated in Snail^6SA^ cells

What then could cause the extrusion of Snail^6SA^ cells? Contractile stress at E-cadherin junctions has been implicated in the extrusion of both oncogene-transfected and apoptotic cells (Wu, Gomez et al. 2014, Michael, Meiring et al. 2016). Accordingly, we interrogated our transcriptomic profiles for changes in potential regulators of the actomyosin cytoskeleton. Interestingly, we found that RhoA mRNA levels were moderately elevated in Snail^6SA^ (Figure 3A), an observation that was confirmed by quantitative PCR (Figure S2B) and by immunoblotting for total RhoA protein (Figure 3D and E). This suggested that Snail^6SA^ might enhance the expression of RhoA. However, RhoA signals to the actomyosin cytoskeleton when it is bound to GTP (Jaffe and Hall 2005). To test if the increased expression of RhoA was accompanied by its increased activity in the cells, we isolated GTP-RhoA using Rhotekin-RBD pull-down assays. These showed that overall GTP-RhoA levels were increased in Snail^6SA^ cells compared with controls (Figure 3B and C). Thus, Snail signalling stimulated both the expression and activation of RhoA in cells.

**Figure 3.**
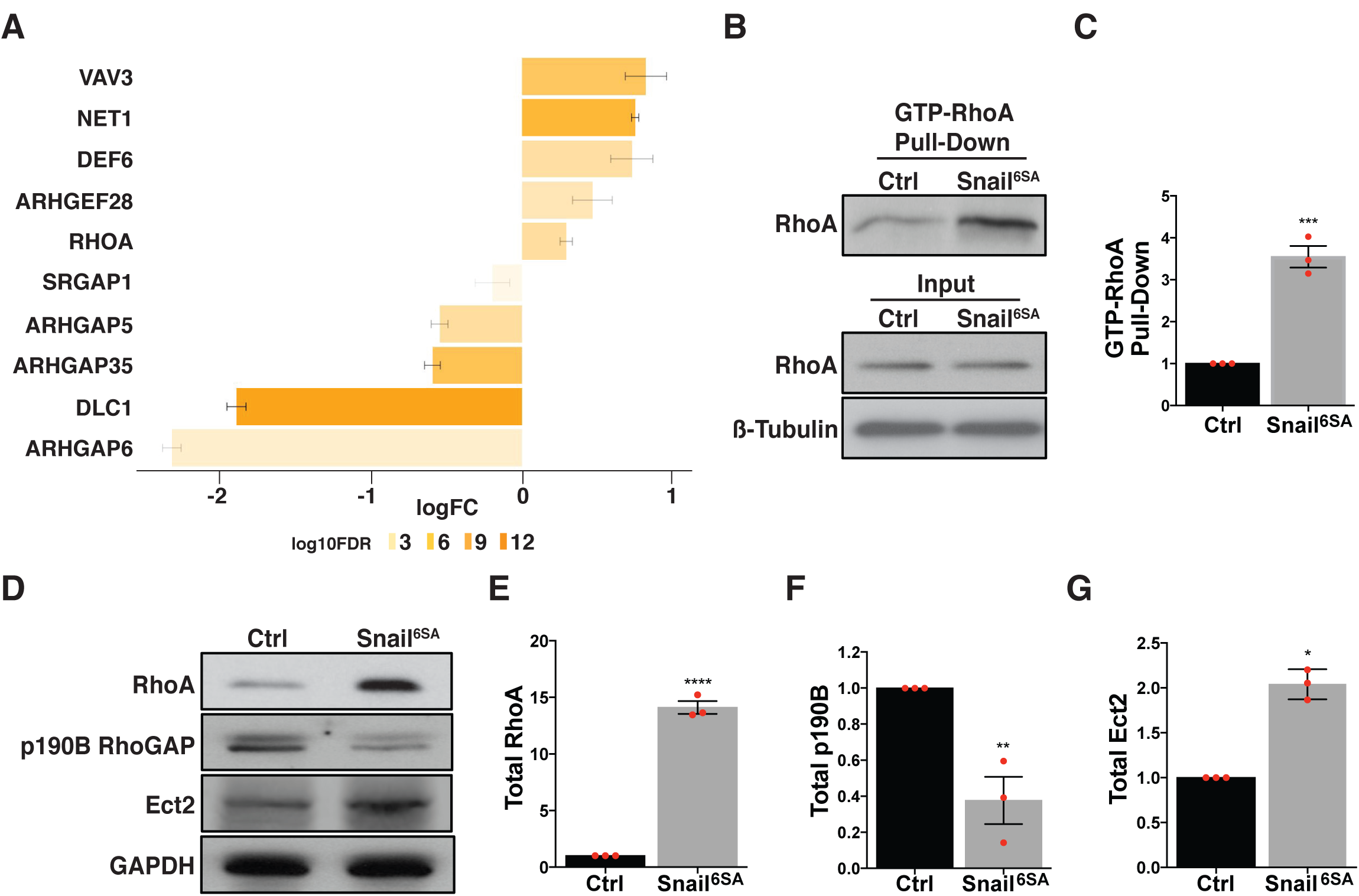
RhoA contractile signaling pathway is hyper-activated in Snail^6SA^ cells. **A)** RNA-seq analysis of regulators of RhoA pathway that are differentially expressed in mCherry^NLS^ versus Snail^6SA^ cells. The length of the bar indicates the fold change of the gene’s expression, while darker colours indicate more strongly significant differential expression. All genes in this plot are significant with a FDR corrected p value <0.001. **B)** Immunoblots of pulled-down active RhoA (GTP-RhoA) **(Top)**, and total RhoA and ß- Tubulin **(bottom)** in cells stably expressing mCherry^NLS^ (Ctrl) or Snail^6SA^. **C)** Quantification of pulled-down GTP-RhoA bands in **(B)**. **D)** Immunoblots of total RhoA, p190BRhoGAP, Ect2 and GAPDH in MCF-7 cells stably expressing mCherry^NLS^ (Ctrl) or Snail^6SA^. **E - G)** Quantification of total RhoA **(E)**, p190BGAP **(F)** and Ect2 **(G)** immunoblots in **(D)**. All mCherry^NLS^ (Ctrl) and Snail^6SA^ cells were treated with 3μg/mL of doxycycline for 48 hours. Data represent means ± S.E.M (*p < 0.05, **p < 0.01, ***p < 0.001) and n = 3 independent experiments; Students t-test (C, E, F, G).

RhoA activity is controlled by the balance between guanine nucleotide exchange factors (GEFs), which promote its active GTP-loaded state, and inhibitory GTPase-activating proteins (GAPs) (Jaffe and Hall 2005). We therefore asked if transcriptional regulation by Snail^6SA^ might affect these classes of proteins. For this, we tested if genes from the curated gene set of the RhoA regulatory pathway were differentially expressed in our cell lines. This revealed that mRNA levels for the RhoA GEFs, NET1, DEF6 and ARHGEF28, were increased in Snail^6SA^ cells compared with controls. Conversely, transcript levels for the RhoA GAPs, DLC1, SRGAP1, ARHGAP5 (p190BRhoGAP) and 6, were reduced upon Snail^6SA^ expression (Fig 3A). We validated some of these changes by Q-PCR and western analysis, choosing candidates for which effective antibodies were available. Together, these experiments corroborated that both mRNA and cellular protein levels of the RhoGEF Ect2 was elevated in Snail^6SA^ cells compared with controls (Figure 3D, G and S2B). Conversely, mRNA and protein levels of p190B RhoGAP were reduced in Snail^6SA^ cells (Figure 3D, F and S2B). Overall, this suggested that transcriptional regulation by Snail^6SA^ could stimulate RhoA signalling both by up-regulating RhoA itself and by altering the GEF:GAP balance to promote its activation.

### Hyper-activated RhoA signals cause elevated tensile forces at E-cadherin junctions in Snail^6SA^ cells

Adherens junctions (AJ) are a prominent site of RhoA signalling in MCF-7 cells. We therefore asked if the increase in overall RhoA activity that we had observed in Snail^6SA^ cells was apparent at their apical adherens junctions. To test this, we used AHPH, a location biosensor for GTP-RhoA (Priya, Gomez et al. 2015, Liang, Budnar et al. 2017), which demonstrated that levels of active RhoA were increased at the AJ and at apical regions in Snail^6SA^ cells compared with controls (Figure 4A - C). Furthermore, a FRET-based RhoA sensor also showed increased activity at the AJ of Snail^6SA^ cells (Figure 4D and E). As this activity sensor reflects the balance between RhoA activators and inactivators, this suggested that the network changes seen in our transcriptomic analysis might also affect the expression of GEFs and GAPs at junctions. Indeed, levels of the RhoGEF Ect2 were elevated at junctions (Figure S3A and B), while levels of p190B RhoGAP were repressed, in Snail^6SA^ cells compared with controls (Figure S3A and C). Thus, we showed that Snail^6SA^ expression stimulates RhoA signalling at the apical adherens junctions.

**Figure 4.**
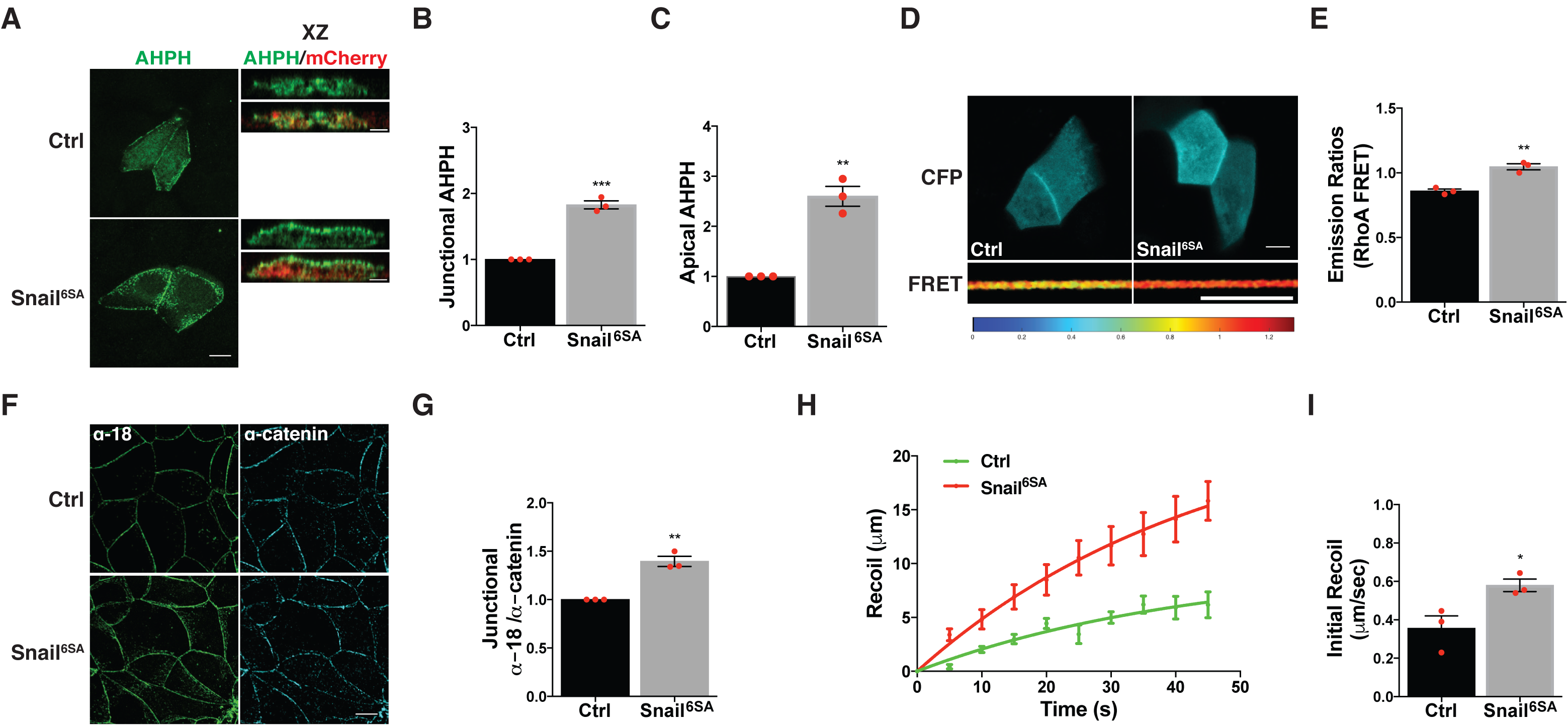
Hyper-activated RhoA signals cause elevated tensile forces at E-cadherin junctions in Snail^6SA^ cells. **A) (Left)** Representative XY images of mCherry^NLS^ (Ctrl) or Snail^6SA^ cells transfected with GFP-AHPH and immunostained with mCherry and GFP. **(Right)** Representative XZ images of cells from the left showing medio-apical localisation of GFP-AHPH, particularly in Snail^6SA^ cells. **B)** Quantification of junctional GFP-AHPH in mCherry^NLS^ (Ctrl) or Snail^6SA^ cells from XY images in **(A, Left).** **C)** Quantification of medio-apical GFP-AHPH in mCherry^NLS^ (Ctrl) or Snail^6SA^ cells from XY images in **(A, Right)**. **D)** Representative live images of CFP **(top)** and RhoA-FRET **(bottom)** at the adherens junctions of mCherry^NLS^ (Ctrl) or Snail^6SA^ cells transfected with pTriEX-RhoA biosensor construct for 12 hours. **E)** Average FRET emission calculation of junctional RhoA-FRET using FRET emission fluorescence as a ratio of doner emission (CFP) in **(D)**. **F)** Representative images of mCherry^NLS^ (Ctrl) or Snail^6SA^ cells immunostained with α-18 and α-catenin antibodies as a proxy for junctional tension. **G)** Junctional fluorescence intensity of α-18 and α-catenin in mCherry^NLS^ (Ctrl) or Snail^6SA^ cells from XY images in **(F)** were quantified as a ratio (α-18:α-catenin). **H, I)** Vertex displacement **(H)** and initial recoil velocity **(I)** of E-cadherin junctions in mCherry^NLS^ (Ctrl) or Snail^6SA^ cells from two-photon laser ablation. All mCherry^NLS^ (Ctrl) and Snail^6SA^ cells were treated with 3μg/mL of doxycycline for 48 hours. Data represent means ± S.E.M (*p < 0.05, **p < 0.01, ***p < 0.001) and n = 3 independent experiments; Students t-test (B, C, E, G, I). Scale bars in XY and XZ images represent 10 μm and 5 μm respectively.

We then asked if this increase in RhoA activity affected the actomyosin cytoskeleton at the AJ. Immunofluorescence staining revealed that cortical levels of F-actin and Myosin II (NMIIA) were increased at the junctions of Snail^6SA^ cells (Figure S3A, D and E), while lacking an increase in total NMIIA expression (Figure S2C and D), suggesting that Snail^6SA^ expression influenced NMIIA localisation rather than its overall expression. This was accompanied by increased junctional accumulation of phosphorylated myosin regulatory light chain (ppMLC), an index of activated Myosin (Figure S3A and F). This implied that actomyosin contractility was increased at the AJ, an effect that would be predicted to increase tensile forces at junctions. To test this, we first stained for a tension-sensitive epitope of α-catenin (α-18 mAb) (Yonemura, Wada et al. 2010). Junctional α-18 mAb staining was increased by expression of Snail^6SA^, evident of increased molecular tension across the cadherin-catenin complex (Figure 4F and G). This was corroborated by measuring the recoil of E-cadherin-GFP junctions after they were cut by laser ablation. Initial recoil speed was elevated in Snail^6SA^ cells compared with controls (Figure 4H and I). However, Snail^6SA^ did not apparently alter viscous drag at junctions, as inferred from the rate constants (*k*-values) of relaxation (Figure S3G). This suggested that the increase in initial recoil velocity principally reflected an increase in junctional tension. Together, these findings imply that Snail expression induces a transcriptional network that upregulates cellular contractility via RhoA.

### Hyper-contractility of Snail^6SA^ cells promotes their apical extrusion amidst normal epithelial cells

The transcriptional network that we have identified had the potential to upregulate RhoA signalling in a cell-autonomous fashion. We then tested if this applied when Snail^6SA^ was mosaically expressed in monolayers. First, we used AHPH to characterize the distribution of GTP-RhoA in Snail^6SA^ cells that were surrounded by wild-type cells. Compared with control GFP-transfected cells, Snail^6SA^ cells showed increased cortical levels of AHPH (Figure 5A and B). This was evident at cell-cell junctions, associated with increased junctional RhoA and Ect2 GEF, and a concomitant decrease in p190B RhoGAP (Figure S4A – D). This suggested that Snail^6SA^ could drive RhoA activation even when neighbour cells were wild-type. Interestingly, there was also an increased accumulation of cortical AHPH at the extra-junctional apical domain in these cells (Figure 5A and C), as we had observed in Snail^6SA^ monolayers (Figure 4A and C). This implied that up-regulation of RhoA was not confined to cell-cell junctions, an inference that was consistent with the diverse range of RhoA regulators that were modulated in Snail^6SA^ cells.

**Figure 5.**
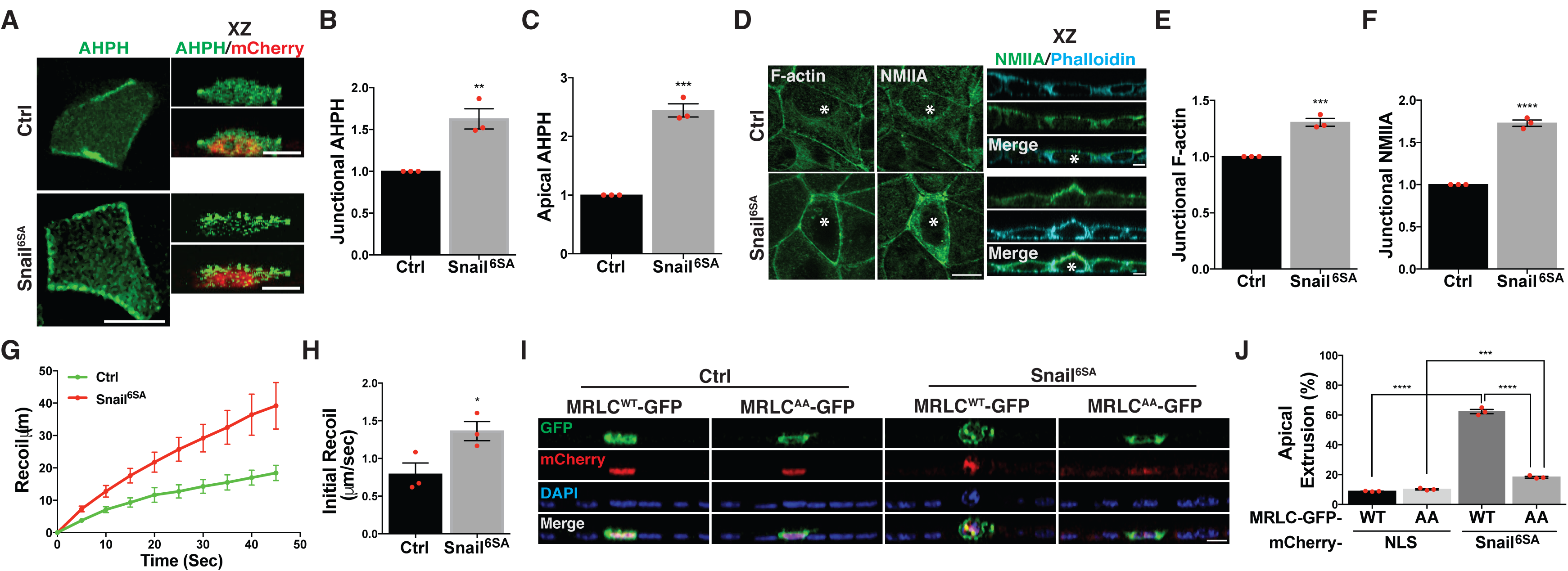
Hyper-contractility of Snail^6SA^ cells promotes their apical extrusion amidst normal epithelial cells. **A) (Left)** Representative XY images of GFP-AHPH transfected mCherry^NLS^ (Ctrl) or Snail^6SA^ cell surrounded by wild type MCF-7 cells and immunostained with mCherry and GFP. **(Right)** Representative XZ images of cells from the left showing medio-apical localisation of GFP-AHPH. **B)** Quantification of junctional GFP-AHPH fluorescence intensity in mCherry^NLS^ or Snail^6SA^ cells at the cell-cell contacts (heterologous interface) between wild type MCF-7 and mCherry^NLS^ (Ctrl) or Snail^6SA^ cells from XY images in **(A, Left)**. **C)** Quantification of medio-apical GFP-AHPH fluorescence intensity in mCherry^NLS^ (Ctrl) or Snail^6SA^ cells from XY images in **(A, Right)**. **D) (Left)** Representative XY images of mCherry^NLS^ (Ctrl) or Snail^6SA^ cells **(asterisk)** in a mosaic environment immunostained with Phalloidin (F-actin) and NMIIA. **(Right)** Still XZ images of cells from **(Left)**, showing accumulation of NMIIA and Phalloidin (F-actin) at the heterologous interface between wild type MCF-7 and mCherry^NLS^ or Snail^6SA^ cells **(asterix)**. **E, F)** Quantification of junctional Phalloidin (F-actin) **(E)** and NMIIA **(F)** fluorescence intensity at the heterologous interface between mCherry^NLS^ (Ctrl) or Snail^6SA^ cells and wild type MCF- 7 cells from XY images in **(D, Left)**. **G, H)** Vertex displacement **(G)** and initial recoil **(H)** of E-cadherin junctions at the heterologous interface between mCherry^NLS^ (Ctrl) or Snail^6SA^ cells and wild type MCF-7 cells from two-photon laser ablation. **I)** Effect of reduced cellular and junctional contractility in the apical extrusion of Snail^6SA^ cells through MRLC^AA^-GFP overexpression. Representative XZ images of wild type MCF-7 cells mixed with either mCherry^NLS^ (Ctrl) or Snail^6SA^ cells stably expressing either MRLC^WT^-GFP or MRLC^AA^-GFP and immunostained with mCherry and GFP. **J)** Quantification of cells in **(I)** undergoing apical extrusion. All mCherry^NLS^ (Ctrl) and Snail^6SA^ cells were treated with 3μg/mL of doxycycline for 48 hours. Data represent means ± S.E.M (*p < 0.05, **p < 0.01, ***p < 0.001) and n = 3 independent experiments; Students t-test (B, C, E, F, H, J). Scale bars in XY and XZ images represent 10 μm and 5 μm respectively.

Immunofluorescence further revealed that cortical levels of F-actin and Myosin II were increased at the apical junctions between Snail^6SA^ cells and their WT neighbours in these mosaic cultures (Figure 5D - F). This was accompanied by evidence of increased mechanical tension at these heterologous junctions, which showed increased staining with the α-18 mAb (Figure S4E and F) and increased initial recoil after laser ablation (Figure 5G and H). Thus, the cell-autonomous up-regulation of RhoA by Snail^6SA^ stimulated junctional actomyosin to enhance contractile tension.

Based on these findings, we hypothesized that the increased contractility in Snail^6SA^ cells might be responsible for their apical extrusion. Consistent with this, we found that depletion of either RhoA or NMIIA throughout the epithelium led to a reduction in the apical extrusion of mosaically-expressed Snail^6SA^ cells (Figure S4G and H). Then, to specifically test if enhanced contractility in the Snail^6SA^ cells was driving extrusion, we expressed a myosin regulatory light chain transgene (MRLC^AA^) in these cells. MRLC^AA^ is mutated for the residues (Thr19, Ser20) whose phosphorylation is necessary to activate NMII. The over-expression of MRLC^AA^ transgene in parental MCF-7 and Snail^6SA^ cells reduced the junctional recoil after laser ablation (Figure S4I – K, M - O), confirming its usefulness to dampen both, contractility and junctional tension. Furthermore, we failed to detect any phosphorylation of MRLC^AA^ at the serine-19 (S19) site, establishing the dependence of MRLC phosphorylation on contractility (Figure S4L). Then, we co-expressed MRLC^AA^ with Snail^6SA^ and mixed these cells with WT cells. Strikingly, under these mosaic conditions we found that MRLC^AA^ reduced the apical extrusion of Snail^6SA^ cells by ∼ 3-fold, compared with Snail^6SA^ cells where wild-type MRLC was expressed (Figure 5I and J). This implied that increased contractility induced in Snail^6SA^ cells was necessary for their apical extrusion.

### Snail^6SA^ induces loss of basal adhesion and promotes delamination

We then sought to understand the cause of delamination when Snail^6SA^ was expressed in large groups of cells. To test if upregulation of contractility was sufficient to explain this phenomenon, we expressed a phosphomimetic MRLC (MRLC^DD^), which imparts elevated tension at apical adherens junctions (Leerberg, Gomez et al. 2014), to mimic the effects of increasing contractility in otherwise WT cells. However, delamination was not enhanced in MRLC^DD^ monolayers (Figure 6A and B). This suggested that enhanced contractility alone did not account for the delamination seen in Snail^6SA^ monolayers.

**Figure 6.**
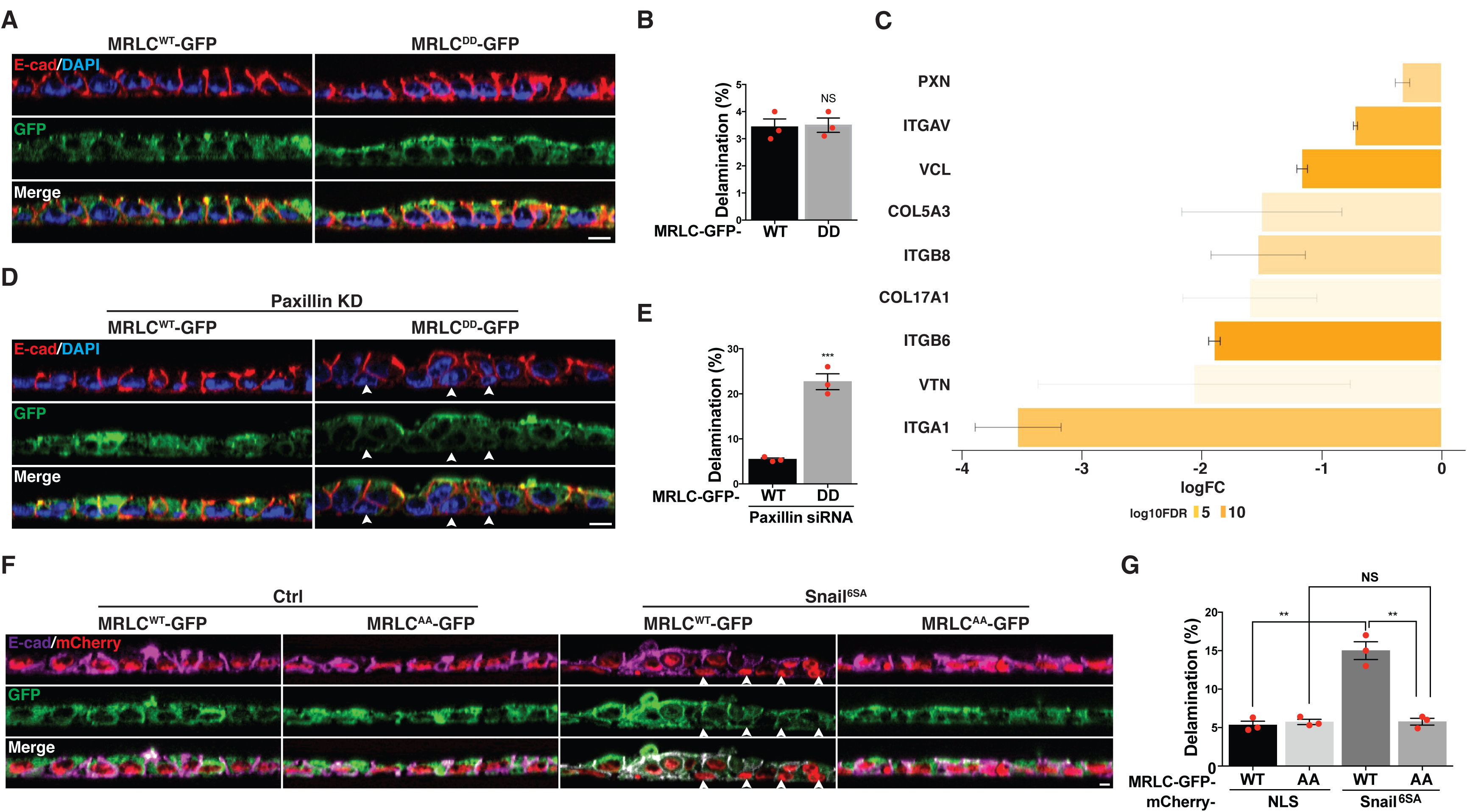
Snail^6SA^ induces loss of basal adhesion and promotes delamination. **A, B)** Representative XZ images **(A)** and quantification **(B)** of delaminated MCF-7 cells expressing either MRLC^WT^-GFP or MRLC^DD^-GFP immunostained with E-cadherin and GFP. **C)** RNA-seq analysis of genes associated with Gene Ontology category ECM matrix adhesion, in mCherry^NLS^ (Ctrl) or Snail^6SA^ stable cells. **D, E)** Representative XZ images **(D)** and quantification **(E)** of delaminated MCF-7 cells expressing either MRLC^WT^-GFP or MRLC^DD^-GFP cells after paxillin knockdown, immunostained with E-cadherin and GFP. **F, G)** Representative XZ images **(F)** and quantification **(G)** of delaminated mCherry^NLS^ (Ctrl) or Snail^6SA^ cells expressing either MRLC^WT^-GFP or MRLC^AA^-GFP immunostained with E- cadherin, mCherry and GFP. Arrows indicate delaminated cells. All mCherry^NLS^ (Ctrl) and Snail^6SA^ cells were treated with 3μg/mL of doxycycline for 48 hours. Data represent means ± S.E.M (*p < 0.05, **p < 0.01, ***p < 0.001 and ns = not significant) and n = 3 independent experiments; Students t-test (B, E, G). Scale bars in XZ images represent 5 μm.

Epithelial cells exhibit strong adhesion to secreted matrix that is required for the polarized and columnar architecture of these cells (Mao and Baum 2015). We therefore hypothesized that Snail^6SA^ potentially affects cell-ECM adhesion leading to delamination of these cells. To test this, we used the gene-sets relating to extra-cellular matrix reorganization, basal lamina and cell matrix adhesion from Reactome and GO databases to assess the enrichment of the DE genes. We observed significant downregulation of expression of genes related to cell-ECM adhesion and extracellular matrix (e.g., Paxillin, Zyxin, Integrins, collagen and laminin), and upregulation of transcripts of genes influencing matrix remodelling (e.g., Matrix metalloproteinases - MMP9, MMP25) (Figure 6C and S5A and B). Immunoblotting of pFAK, vinculin and paxillin and immunostaining of paxillin confirmed the loss of cell-ECM adhesion in Snail^6SA^ monolayers (Figure S6A - F). It was interesting to note that Snail^6SA^ cells also showed elevated number of invadopodia per cell (Figure S6G - I). The loss of cell-ECM adhesion along with matrix remodelling could thus be a potential cause of invadopodia in Snail^6SA^ cells, as reported earlier in other contexts (Chan, Cortesio et al. 2009). These findings imply that transcriptional activities of Snail induces loss of cell-ECM adhesion along with increased matrix remodelling.

To test if loss of cell-ECM adhesion can trigger basal delamination, we depleted paxillin in MCF-7 monolayers by siRNA (Figure S6J). Loss of paxillin led to a moderate increase in basal cell delamination in MCF-7 monolayers (Figure S6K and L). This suggested that the strength of cell-ECM contributed to delamination but may not be solely responsible for delamination in Snail^6SA^ monolayers (Figure 1F and G) due to a discrepancy in the delamination rates. We then hypothesized that loss of cell-ECM adhesion coupled with elevated contractility, as in the case of Snail^6SA^ expression, could further promote the delamination process. To test this, we expressed MRLC^DD^ in cells depleted of paxillin to simultaneously induce hyper-contractility and reduce cell-ECM adhesions. The increase in cortical contractility coupled with loss of cell-ECM adhesion led to a drastic enhancement in delamination of cells in MCF-7 monolayers (Figure 6D and E), similar to that observed in Snail^6SA^ monolayers (Figure 1F and G and Supplemental Movie 2). To further confirm this, we dampened contractility in Snail^6SA^ monolayers by expression of MRLC^AA^. Expression of MRLC^AA^ in Snail^6SA^ monolayers suppressed delamination to a minimal level as seen in control conditions (Figure 6F and G). These findings indicate a direct role for cortical contractility coupled to loss of basal adhesion for delamination in Snail^6SA^ monolayers.

## Discussion

Epithelial tissues elicit a protective response to insults by expelling aberrant and damaged cells apically, a process known as apical extrusion (Hogan, Dupre-Crochet et al. 2009, Slattum, Mcgee et al. 2009, Michael, Meiring et al. 2016). These insults can be lethal (apoptotic inducing), or pathological (oncogenic transformation), which then triggers the extrusion of such cells. While the morphogenetic factor, Snail, is a major driver of epithelial morphogenesis during development, its aberrant expression in adults contributes to processes such as invasion and migration during malignant disease (Alberga, Boulay et al. 1991, Nieto 2002, Peinado, Olmeda et al. 2007). Here, we show that mosaic expression of a stable Snail transgene (Snail^6SA^) in an epithelium led to the apical extrusion of these transgene-expressing cells. Apical extrusion arose in response to the elevated contractility of the mosaic Snail^6SA^ cells that surprisingly did not concomitantly involve EMT. Thus, it closely resembles the extrusion response to epithelial apoptosis, where enhanced contractility in the apoptotic cell stimulates its neighbours to drive apical extrusion (Gagliardi, Somale et al. 2018) (Michael, Meiring et al. 2016). However, the extrusion of Snail^6SA^ cells was not accompanied by apoptosis, suggesting that the principal stimulus may have been mechanical.

While its ability to promote EMT is the best-understood way for Snail to exert its morphogenetic effects (Wu and McClay 2007, Theveneau and Mayor 2012), Snail expression could also contribute to local contractions of the actomyosin network during development, as in the context of apical constriction (Martin, Kaschube et al. 2009, Martin, Gelbart et al. 2010, Weng and Wieschaus 2016). However, the mechanisms responsible for this latter effect are poorly characterized. Here, we find that this can be explained by the modulation of a transcriptional network leading to the stimulation of RhoA signalling in the MCF-7 epithelial cell system. Surprisingly, the precise morphogenetic consequences of this pathway depended critically on the extent to which Snail was expressed within monolayers. Although expression of Snail^6SA^ in small groups of cells (i.e. <3) led to their apical extrusion, the integrity of the monolayers remained largely intact when Snail^6SA^ was expressed throughout the monolayers. Cortical actomyosin and contractility was upregulated, as evident by an increase in mechanical tension at the AJ. However, monolayers largely retained epithelial organization. This is reminiscent of what has been observed when oncogenes, such as Ras and Src, are ubiquitously expressed in otherwise differentiated epithelial monolayers (Hogan, Dupre-Crochet et al. 2009, Kajita, Hogan et al. 2010). We hypothesize that a key determinant of outcome was the balance of contractile forces at the tissue level. Mosaic expression of Snail^6SA^ would be predicted to cause a local upregulation of contractility whereas its ubiquitous expression would be expected to lead to an overall increase in tissue tension that was balanced throughout the monolayer. A role for local imbalances in force is further supported by the observation that extrusion was limited to small clusters of cells (< 3 cells). E-cadherin-based adherens junctions possess mechanotransduction pathways that can respond to mechanical stresses (Acharya, Nestor-Bergmann et al. 2018). However, whether cell-autonomous activation of contractility is sufficient to drive apical extrusion of Snail^6SA^ cells or also requires an active response of the neighbour cells remains to be determined.

Although overall epithelial integrity was preserved when Snail^6SA^ was expressed ubiquitously, we noted that under these circumstances some cells were basally delaminated. This appeared to be a stochastic process, as only a minority of the transgene-expressing cells delaminated. This contrasts with what we saw when Snail^6SA^ was expressed mosaically, where the majority of the transfected cells were apically extruded. Here we found that upregulation of contractility was also necessary but not sufficient for basal delamination. Whereas apical extrusion can be elicited by mosaic upregulation of contractility alone, increased junctional tension alone, through ubiquitous expression of MRLC^DD^, did not promote delamination. This finding was intriguing as hyper-contractility was sufficient to trigger delamination in developing embryos (Liu and Jessell 1998). Instead, our data indicate that basal delamination requires a combination of enhanced contractility together with changes in cell-ECM interactions. Our transcriptomic analysis revealed that Snail^6SA^ expression modulated a suite of genes that are predicted to regulate integrin-based adhesion and enhance turnover of the ECM. Consistent with this, we found that focal adhesions were less apparent in Snail^6SA^ cells and paxillin RNAi modestly increased basal delamination. However, it was the combination of paxillin RNAi and MRLC^DD^ that most strikingly enhanced delamination, thus implicating a critical role for both cortical contractility and basal adhesion in this process. This notion was further reinforced by the observation that basal delamination in Snail^6SA^ monolayers was suppressed when cortical contractility was inhibited by expression of MRLC^AA^. Together these findings indicate a critical role for Snail to transcriptionally control contractility via the RhoA pathway and also basal adhesion to drive expulsion of cells from the epithelia that is independent of its canonical role of EMT. While the role of cell-ECM interactions in inducing basal delamination remains unclear, cross-talks between molecules at apical junctions and sites of basal adhesion could be a potential trigger. The loss of basal adhesion may facilitate reinforcement of apical adherens junctions that could induce a loss of apical surface and the subsequent basal expulsion of these cells.

In conclusion, we propose that Snail induced changes in cell contractility and basal adhesion as a mechanism for epithelial cells to basally escape the constraints of tissue environment to initiate invasive mechanisms. These mechanisms could be potentially co-opted by oncogenic cells to escape tissues, despite resistive E-cadherin junctions. While the induction of EMT pathways by Snail family members is possibly dependent on the cellular context, the observed preservation of an epithelial identity to escape the tissue environment potentially uncovers new roles for Snail in the modulation of epithelial cell behaviour, as similarly identified during the dissemination of Twist cells from the mammary epithelia (Shamir, Pappalardo et al. 2014).

## Acknowledgements

We would like to thank our laboratory colleagues, past and present, for their immense support and advise. This work was supported by an Australian Postgraduate Award, to KW, and grants and fellowships from the Queensland Cancer Council (1086587, 112823), and the National Health and Medical Research Council of Australia (1044041, 1136592, 1067405) to AY. MJD is supported by National Breast Cancer Foundation (ECF-14-043 and CG-10-04; funding of the EMPathy Breast Cancer Network) and the Australian Research Council Center of Excellence in Convergent Bio-Nano Science and Technology (project number CE140100036). RJD is supported by NHMRC Fellowship APP1058540.

## Author contributions

KW, ASY, SB conceived the project; KW designed and performed experiments, and analysed the data. SV provided technical assistance for GTP-RhoA pull down assay. MJD and SZ performed the RNA-seq analyses. SB designed experiments, generated the GFP tagged E-cadherin genome edited MCF-7 cells, generated inducible expression constructs, supervised the work and coordinated the project. KW, MD, ASY and SB wrote the paper.

---

**Figure S1, related to Figure 1.**
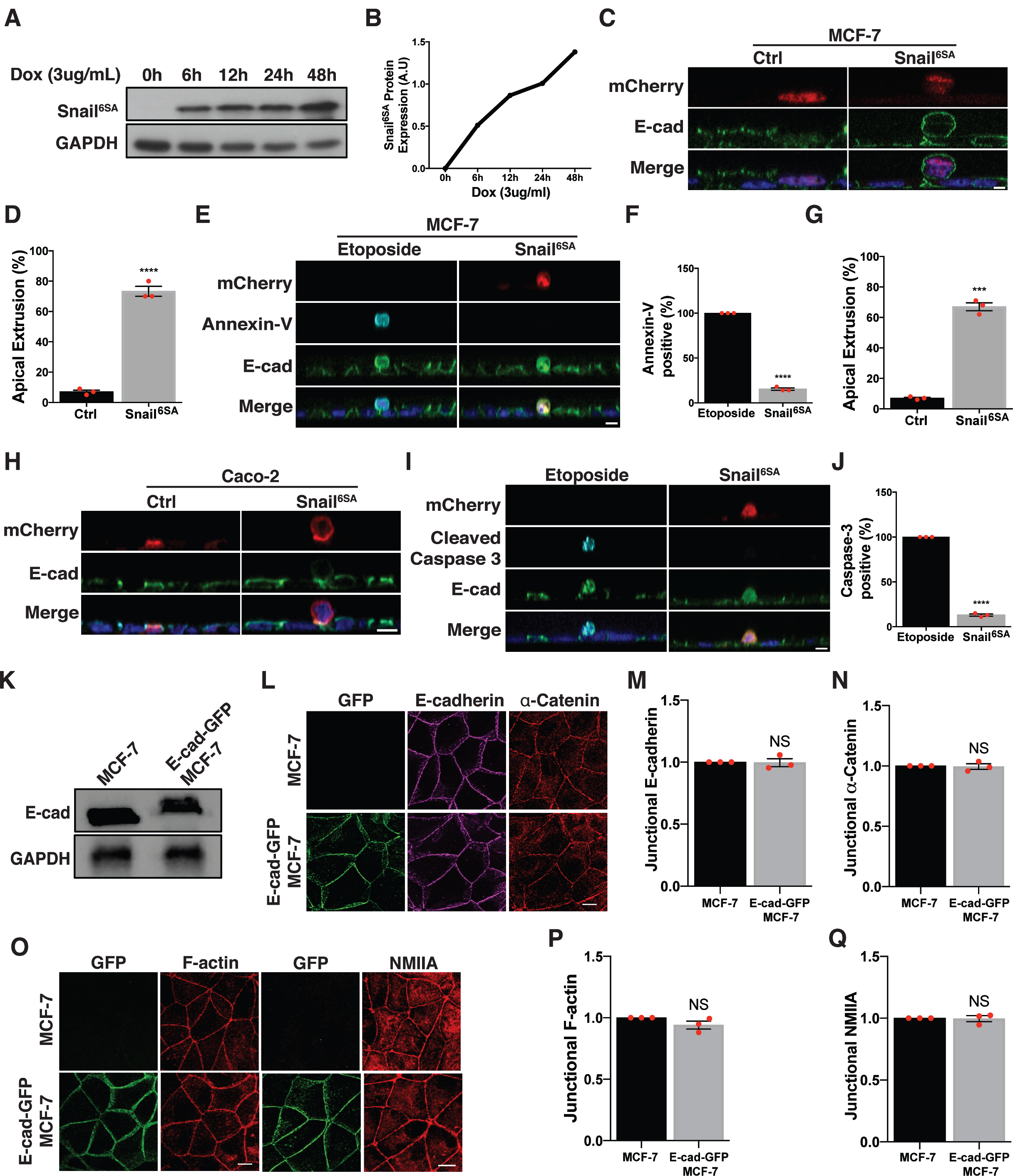
Snail^6SA^ induced apical extrusion is independent of apoptosis and cell type. **A, B)** Immunoblots **(A)** and quantification **(B)** of Snail^6SA^ expression chased at the indicated times upon 3μg/mL of doxycycline treatment. **C, D)** Representative XZ images **(C)** and quantification **(D)** of apically extruded mCherry^NLS^ (Ctrl) or Snail^6SA^ cells when mixed with wild type MCF-7 cells in a mosaic environment. **E, F)** Representative XZ images **(E)** and quantification **(F)** of either apically extruded Snail^6SA^ in a mosaic population or MCF-7 cells treated with etoposide (250nM) immunostained with Annexin-V as an apoptotic marker. **G, H)** Quantification **(G)** and representative XZ images **(H)** of apically extruded Caco-2 cells stably expressing mCherry^NLS^ (Ctrl) or Snail^6SA^ mixed with wild type Caco-2 cells. **I, J)** Representative XZ images **(I)** and quantification **(J)** of either apically extruded Snail^6SA^ Caco-2 cells in a mosaic population or Caco-2 cells treated with etoposide (250nM) immunostained with cleaved caspase-3 as an apoptotic marker. All mCherry^NLS^ (Ctrl) and Snail^6SA^ cells were treated with 3μg/mL of doxycycline for 48 hours, except for **(A)** and **(B)**. **K)** Immunoblots of total E-cadherin and GAPDH in parental MCF-7 cells (MCF-7) or MCF-7 cells in which the endogenous E-cadherin is tagged with GFP (E-cad-GFP-MCF-7). **L)** Representative XY images of parental MCF-7 cells (MCF-7) or E-cad-GFP-MCF-7 cells immunostained with GFP, E-cadherin and α-catenin. **M, N)** Quantification of junctional E-cadherin **(M)** and α-catenin **(N)** fluorescence intensity in parental MCF-7 cells (MCF-7) or E-cad-GFP-MCF-7 cells in **(L)**. **O)** Representative XY images of parental MCF-7 cells (MCF-7) or E-cad-GFP-MCF-7 cells immunostained with GFP, Phalloidin (F-actin) and NMIIA. **P, Q)** Quantification of junctional F-actin **(P)** and NMIIA **(Q)** fluorescence intensity in parental MCF-7 cells (MCF-7) or E-cad-GFP-MCF-7 cells in **(O)**. Data represent means ± S.E.M (*p < 0.05, **p < 0.01, ***p < 0.001 and ns = not significant) and n = 3 independent experiments; Students t-test (D, F, G, J, M, N, P, Q). Scale bars in XY and XZ images represent 10 μm and 5 μm respectively.

**Figure S2, related to Figure 3.**
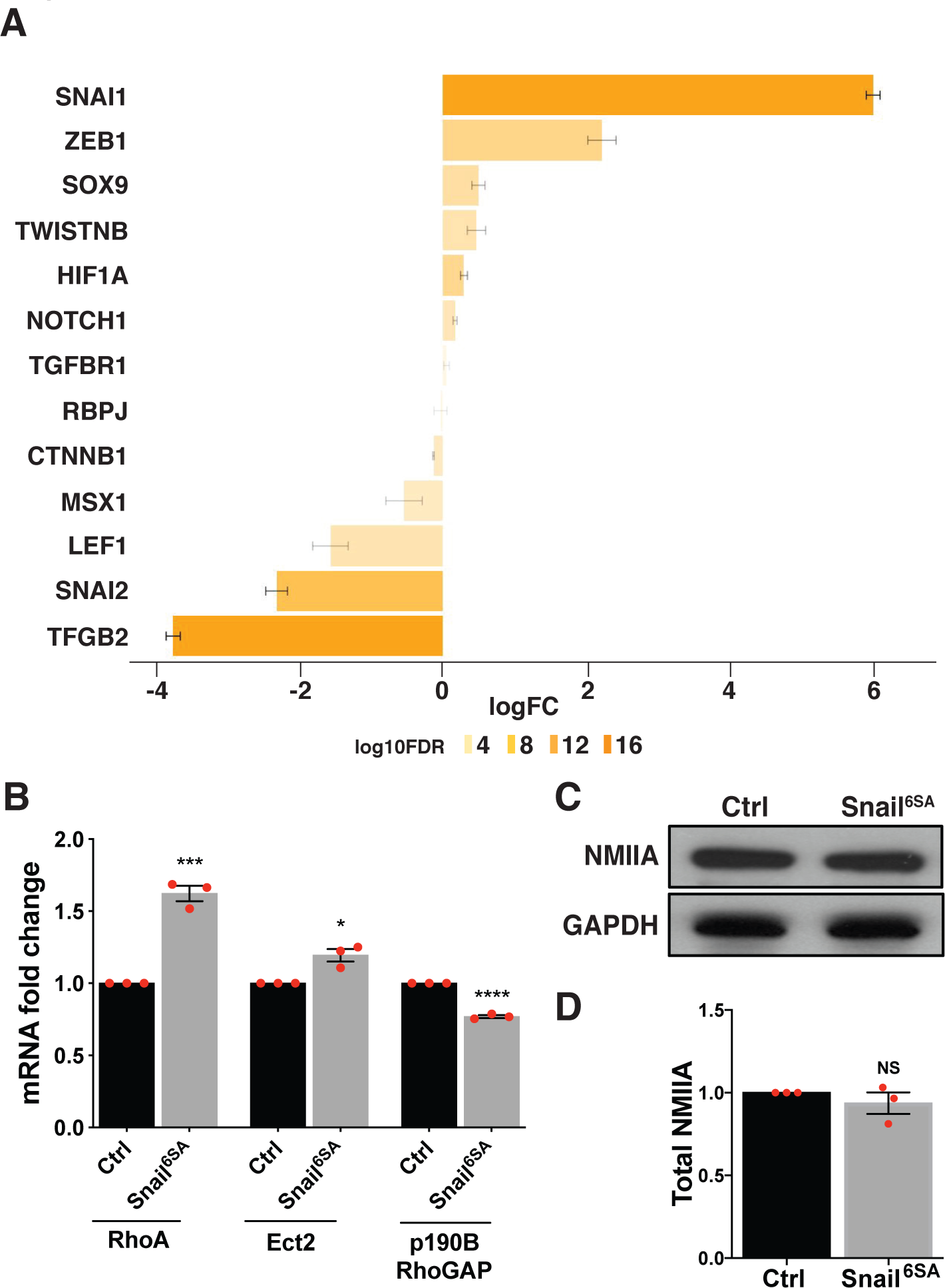
Snail^6SA^ expression regulates RhoA activation. **A)** RNA-seq analysis of transcription factors associated with EMT reprogramming in mCherry^NLS^ (Ctrl) or Snail^6SA^ stable cells. Genes are selected from the union of Gene Ontology categories Transcriptional Regulation and Epithelial Mesenchymal Transition. **B)** mRNA expression of RhoA, Ect2 and p190BRhoGAP in mCherry^NLS^ (Ctrl) or Snail^6SA^ stable cells through RT-qPCR. **C)** Immunoblots of total non-muscle myosin IIA (NMIIA) and GAPDH in MCF-7 cells stably expressing either mCherry^NLS^ (Ctrl) or Snail^6SA^. **D)** Quantification of total NMIIA immunoblot in **(C)**. All mCherry^NLS^ (Ctrl) and Snail^6SA^ cells were treated with 3μg/mL of doxycycline for 48 hours. Data represent means ± S.E.M (*p < 0.05, **p < 0.01, ***p < 0.001 and ns = not significant) and n = 3 independent experiments; Students t-test (B, D).

**Figure S3, related to Figure 3 and 4.**
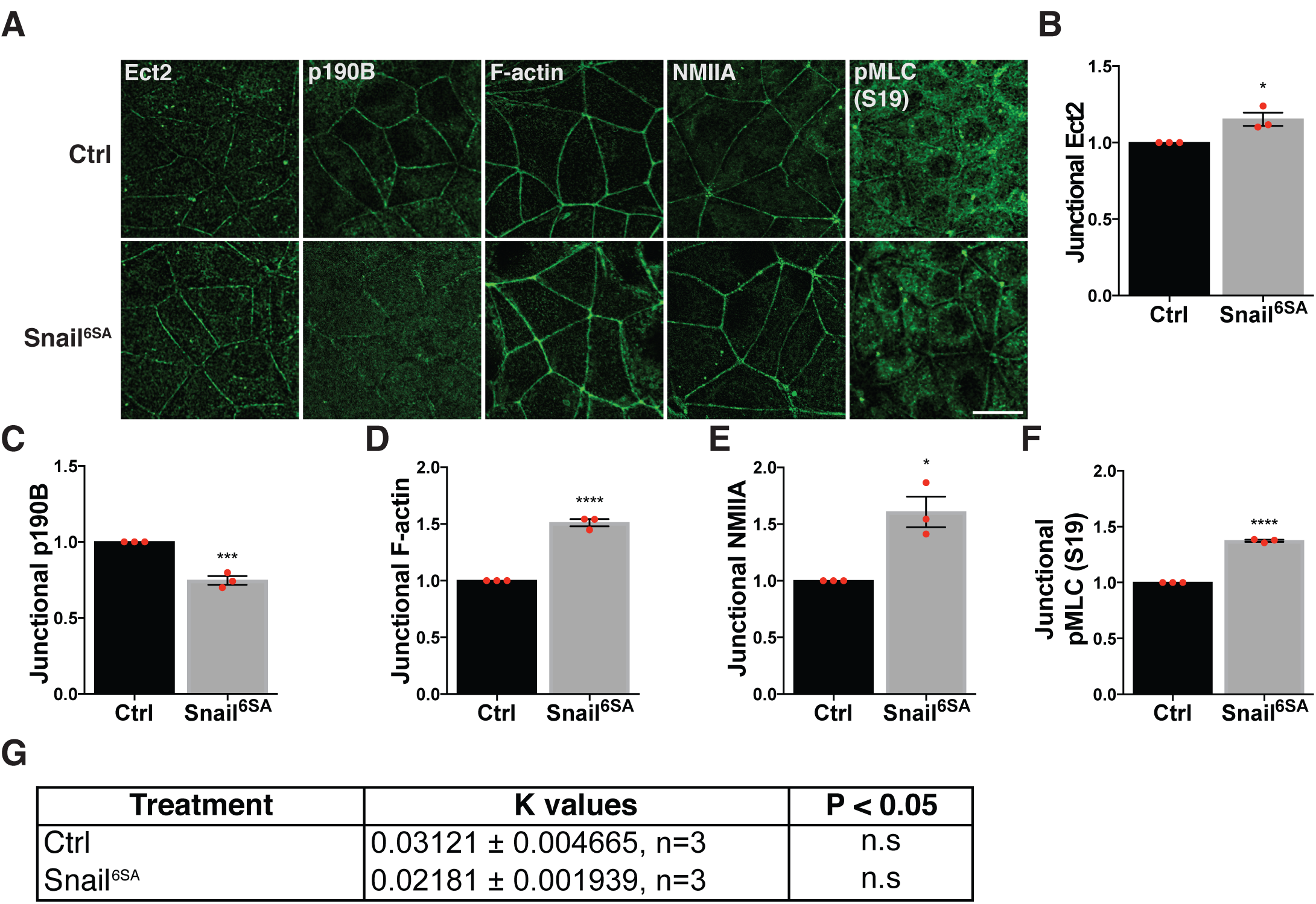
RhoA activation in Snail^6SA^ cells promotes accumulation of active junctional myosin. **A)** Representative XY images of mCherry^NLS^ (Ctrl) or Snail^6SA^ MCF-7 cells immunostained with Ect2, p190BGAP, Phalloidin (F-actin), NMIIA, and phospho-serine19-MLC (pMLC). **B – F)** Quantification of junctional Ect2, p190BGAP, F-actin, NMIIA and pMLC fluorescence intensity from **(A)**. **G)** Assessment of the influence of viscous drag (k-value) on the initial recoils in Fig. 4I. This was achieved by fitting the vertex displacement values from Fig. 4I to a mono-exponential curve to obtain the average rate constant (k) over several experiments. All mCherry^NLS^ (Ctrl) and Snail^6SA^ cells were treated with 3μg/mL of doxycycline for 48 hours. Data represent means ± S.E.M (*p < 0.05, **p < 0.01, ***p < 0.001 and ns = not significant) and n = 3 independent experiments; Students t-test (B, C, D, E, F). Scale bars in XY images represent 10 μm.

**Figure S4, related to Figure 5.**
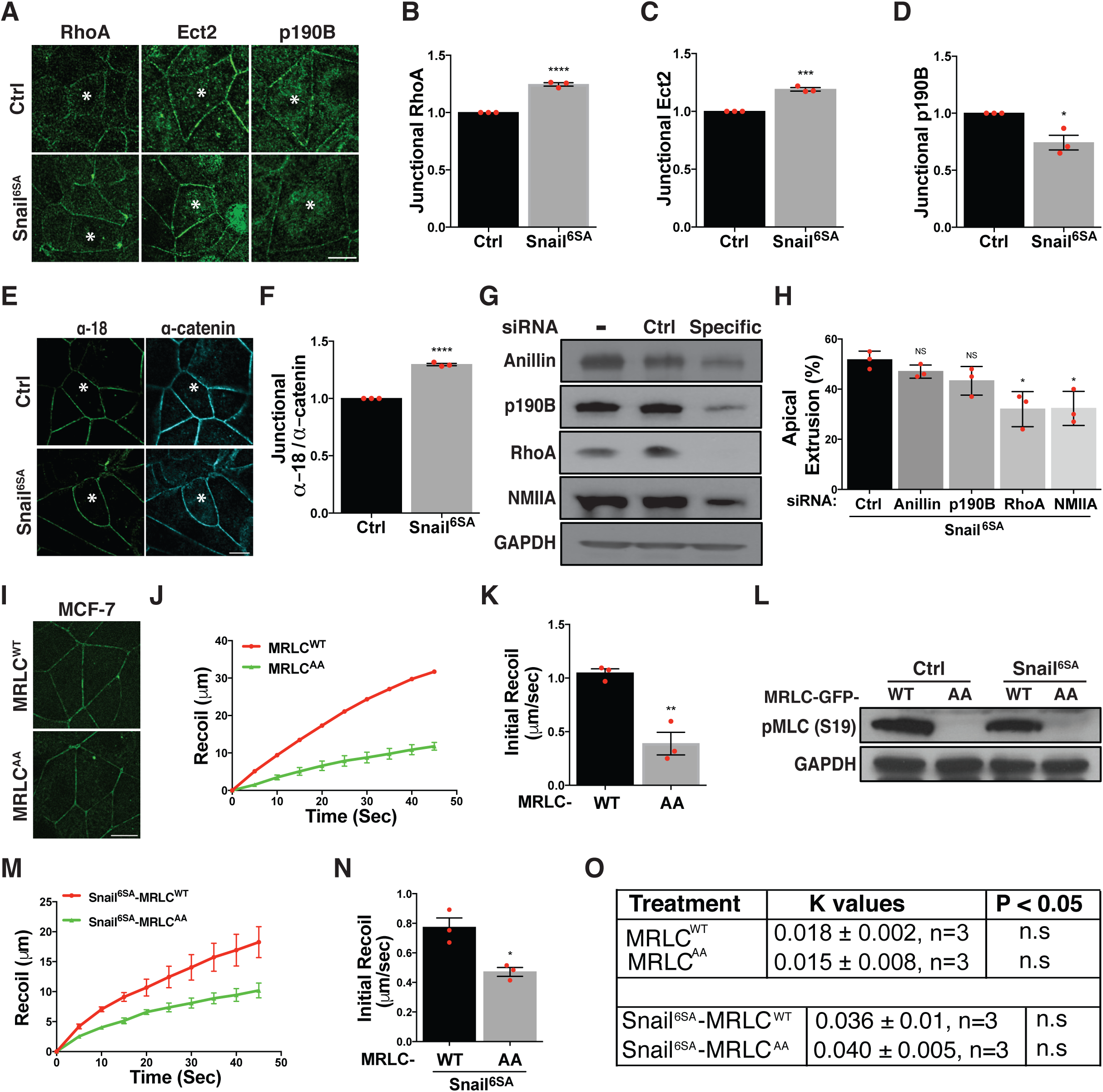
Hyper-activated RhoA signals promote hyper-contractility and apical extrusion of Snail^6SA^ cells. **A)** Representative XY images of MCF-7 cells mixed with either mCherry^NLS^ (Ctrl) or Snail^6SA^ cells **(asterisk)** and immunostained with RhoA, Ect2 and p190BGAP. **B – D)** Quantification of junctional RhoA **(B)**, Ect2 **(C)** and p190BGAP **(D)** fluorescence intensity at the heterologous interface between wild type MCF-7 and mCherry^NLS^ (Ctrl) or Snail^6SA^ cells from (**A)**. **E)** Representative XY images of MCF-7 cells mixed with either mCherry^NLS^ (Ctrl) or Snail^6SA^ cells **(asterix)** and immunostained with α-18 and α-catenin antibodies. **F)** Junctional α-18 and α-catenin fluorescence intensity at the heterologous interface between wild type MCF-7 and mCherry^NLS^ (Ctrl) or Snail^6SA^ cells from **(E)** were quantified as a ratio (α-18:α-catenin). **G)** MCF-7 cells mixed with Snail^6SA^ cells were transfected with control (ctrl), Anilin, p190BGAP, RhoA or NMIIA specific siRNA and immunoblotted for the corresponding siRNA targets. **H)** Quantification of Snail^6SA^ cells from **(G)** undergoing apical extrusion. **I)** Representative live XY images of MCF-7 cells stably expressing either MRLC^WT^-GFP or MRLC^AA^-GFP. **J, K)** Vertex displacement **(J)** and initial recoil velocity (**K)** of adherens junctions in MCF-7 cells expressing either MRLC^WT^-GFP or MRLC^AA^-GFP from **(I)** from two-photon laser ablation. **L)** Immunoblots of pMLC (S19) and GAPDH in mCherry^NLS^ (ctrl) or Snail^6SA^ cells stably expressing either MRLC^WT^-GFP or MRLC^AA^-GFP as a proxy for myosin activation. **M, N)** Vertex displacement **(M)** and initial recoil velocity **(N)** of adherens junctions in Snail^6SA^ cells expressing either MRLC^WT^-GFP or MRLC^AA^-GFP from two-photon laser ablation. **O)** Assessment of the influence of viscous drag (k-value) on the initial recoils in **(K)** and **(N)** respectively. This was achieved by fitting the vertex displacement values from **(K)** and **(N)**, respectively, to a mono-exponential curve to obtain the average rate constant (k) over several experiments. All mCherry^NLS^ (Ctrl) and Snail^6SA^ cells were treated with 3μg/mL of doxycycline for 48 hours, except for **I - K**. Data represent means ± S.E.M (*p < 0.05, **p < 0.01, ***p < 0.001 and ns = not significant) and n = 3 independent experiments; Students t-test (B, C, D, F, H, K, N). Scale bars in XY images represent 10 μm.

**Figure S5, related to Figure 6.**
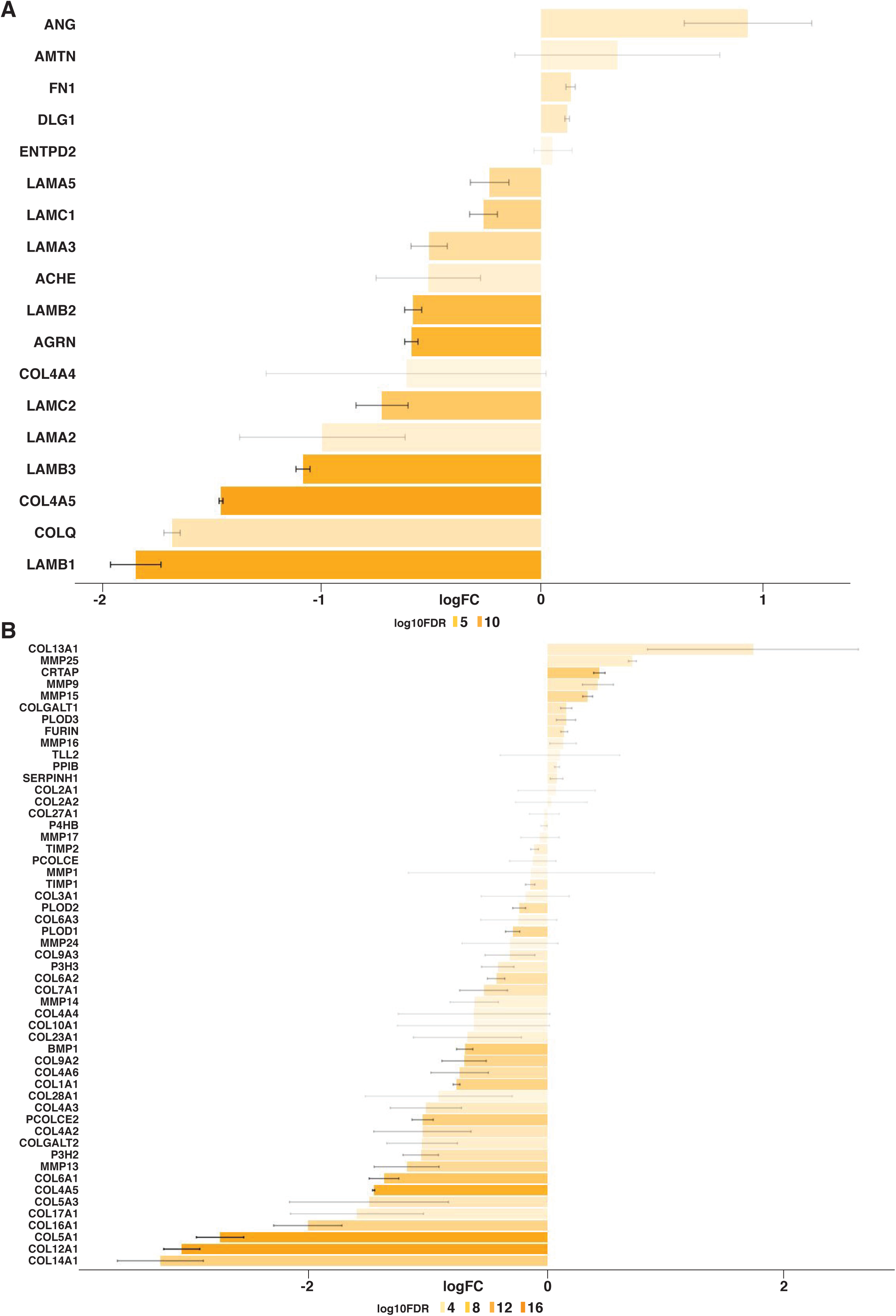
Snail^6SA^ induces changes in basal adhesion and ECM organization. **A, B)** RNA-seq analysis of secreted basal lamina molecules **(A)** and molecules associated with ECM reorganization **(B)** that are differentially expressed in mCherry^NLS^ (Ctrl) versus Snail^6SA^ cells, showing a strong downregulation of genes in these categories. All mCherry^NLS^ (Ctrl) and Snail^6SA^ cells were treated with 3μg/mL of doxycycline for 48 hours.

**Figure S6, related to Figure 6.**
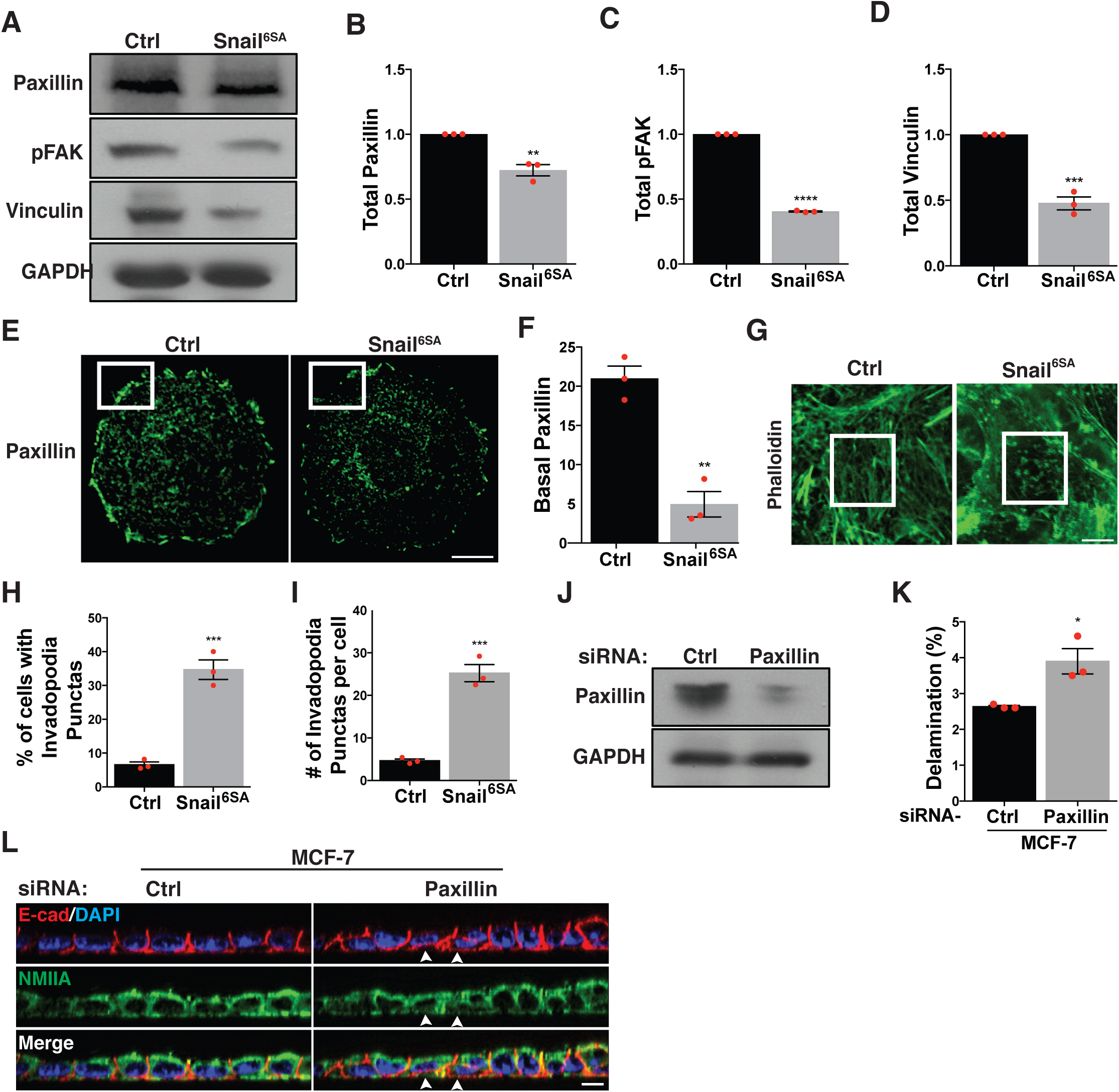
Snail^6SA^ induces loss of basal adhesion and promotes delamination. **A)** Immunoblots of total paxillin, phospho-FAK (pFAK), vinculin and GAPDH in MCF-7 cells stably expressing either mCherry^NLS^ (Ctrl) or Snail^6SA^. **B - D)** Quantification of total paxillin **(B)** pFAK **(C)** and vinculin **(D)** immunoblots in **(A)**. **E)** Representative XY images of single MCF-7 cell stably expressing either mCherry^NLS^ (Ctrl) or Snail^6SA^ immunostained for paxillin. White box indicate paxillin localization at the lamellipodia. **F)** Quantification of paxillin fluorescence intensity at the lamellipodia as indicated by white box in **(E)**. **G)** Representative basal XY images of either mCherry^NLS^ (Ctrl) or Snail^6SA^ cells immunostained for F-actin using phalloidin. White box indicate formation of F-actin invadopodia-like punctas localised beneath the nucleus. **H)** Quantification of mCherry^NLS^ (Ctrl) or Snail^6SA^ cells displaying basal F-actin punctas from **(G)**. **I)** Quantification of the number of basal F-actin punctas in mCherry^NLS^ (Ctrl) or Snail^6SA^ cells from **(G)**. **J)** Immunoblots of paxillin and GAPDH in MCF-7 cells transfected with either control (Ctrl) or paxillin specific siRNA. **K, L)** Quantification **(K)** and representative XZ images **(L)** of MCF-7 cells transfected with either control (Ctrl) or paxillin specific siRNA undergoing delamination, immunostained with E-cadherin and NMIIA. Data represent means ± S.E.M (*p < 0.05, **p < 0.01, ***p < 0.001 and ns = not significant) and n = 3 independent experiments; Students t-test (B, C, D, F, H, I, K). Scale bars in XY and XZ images represent 10 μm and 5 μm respectively.

