## Supplemental Methods for "Snail induces epithelial cell extrusion through transcriptional control of RhoA contractile signaling and cell matrix adhesion"

### Experimental Procedures

#### Cell Culture and transfection

MCF-7 human breast epithelial adenocarcinoma cells were cultured in DMEM complete growth medium (Gibco) supplemented with 10% FBS (Invitrogen), 100 units/mL Penicillin (Gibco) and 100 units/mL Streptomycin (Gibco).

MCF-7 cells stably expressing $mCherry^{NLS}$ or mCherry-$Snail^{6SA}$ were cultured in the same medium as parental MCF-7 cells supplemented with 60 ng/mL of puromycin. MCF-7 cells stably expressing $MRLC^{WT}$-GFP, $MRLC^{AA}$-GFP or $MRLC^{DD}$-GFP were cultured in the same medium as parental MCF-7 cells. Expression of inducible transgenes was achieved by culturing cells with 3μg/mL of doxycycline for at least 48 hours.

Caco-2 human colorectal epithelial carcinoma cells were cultured in RPMI complete growth medium (Gibco) supplemented with 10% FBS (Invitrogen), 100 units/mL Penicillin (Gibco), 100 units/mL Streptomycin (Gibco), 2mM L-glutamine (Gibco) and 1% non-essential amino acids (Gibco).

HEK-293T cells were cultured in the same medium as parental MCF-7 cells. All cells were cultured at 37°C with 5% $CO_2$.

RNAiMax (Invitrogen) and Lipofectamine 2000 (Invitrogen) were used for transfection of siRNA and plasmids respectively, according to the manufacturer's instructions. All analyses of transfected cells were done 48 hours post-transfection, in a confluent monolayer.

#### Plasmids and Generation of Stable Cell Lines

$mCherry^{NLS}$ and mCherry-$Snail^{6SA}$ in pmCherry-C1 (Clontech) were subcloned into the pTripz Tet-ON lentiviral expression vector (GE-Dharmacon), that has been modified to contain a hPGK promoter to drive the expression of rtTA3, using the AgeI and ClaI restriction sites (Table 1).

$MRLC^{WT}$-GFP, $MRLC^{AA}$-GFP and $MRLC^{DD}$-GFP in pmCherry-C1 or pEGFP-C1 (Clontech) were subcloned into the pLL5.0 lentiviral expression vector using the BsrGI and BamHI restriction sites (Table 1). The pLL5.0 lentiviral vector and packaging constructs pMDLg/pRRE, RSV-Rev and pMD.G were gifts from Dr. Jim Bear (UNC Chapel Hill, North Carolina, USA).

To generate the above cell lines, pTripz or pLL5.0 vectors were transfected together with the packaging plasmids into HEK-293T cells using Lipofectamine 2000. Media was replaced 16 hrs post-transfection and the medium-containing virus particles was then collected 72 hrs post-transfection. This medium was then used to infect cells supplemented with 8 µg/mL of polybrene. Cells infected with pTripz viral particles were cultured in media containing 0.5 µg/mL puromycin for 1 week to positively select for transduced cells.

The RhoA location biosensor (GFP-AHPH) is a kind gift from M. Glotzer (University of Chicago, USA) and is described previously (Liang et al., 2017; Priya et al., 2015). To generate a stable cell line, MCF-7 cells were transfected with GFP-AHPH using lipofectamine 2000 and treated with G418 (0.5mg/mL) for selection of positive cell clones. mCherry$^{NLS}$ or Snail$^{6SA}$ expressing cells were transiently transfected with GFP-AHPH using lipofectamine 2000 to analyse for active junctional RhoA.

**RNA isolation, sequencing and qPCR**

MCF-7 cells stably expressing mCherry$^{NLS}$ or mCherry-Snail$^{6SA}$ were treated with 3 µg/mL of doxycycline for 48 hours to induce expression of transgenes. Cells were cultured to 95% confluency and total RNA was extracted using a RNA extraction kit (Sigma) according to the manufacturer's instructions. Purified RNA was enriched for mRNA and library preparation was performed using the TruSeq stranded mRNA kit by the IMB Sequencing Facility (University of Queensland, Institute of Molecular Bioscience). mRNA samples were then sequenced using the NextSeq 75 cycle kit (1 x 75bp) on Illumina NextSeq 500.

For qPCR, RNA was extracted from mCherry$^{NLS}$ or mCherry-Snail$^{6SA}$ expressing cells as described above. 5 µg of purified RNA from each sample was used to convert to cDNA using the SuperScript III First-Strand Synthesis System for RT-PCR (Invitrogen) according to the manufacturer's instructions. Quantification of cDNA samples were performed in a 96 well plate compatible with the ViiA7 Real-Time PCR System (ThermoFisher Scientific). Transcript level of each gene target was assessed in triplicates with specific primers (Table 1). In each replicate well, 4µL of cDNA was added to 0.5 µL each of forward primer and reverse primers (Table 1) and 5 µL of SYBR Green mastermix (Applied Biosystems). The 96-well plate was then briefly vortexed and centrifuged at 1000 rpm for 2 minutes at 4°C. qPCR of the 96 well plate was then performed using the ViiA7 PCR system and the output data was then further analyzed on excel as outlined (Table 2 and 3).

In brief, the CT (cycle threshold) value indicates the number of PCR cycles required for the fluorescent signal of the target to cross the base-line threshold. Thus the CT value obtained from the ViiA7 system is inversely proportional to the amount of mRNA target in the sample. The CT mean value was used as a measurement of relative mRNA expression for each target (Table 2). Relative mRNA levels for each target were then normalized to those of GAPDH in each condition. The fold change of normalized mRNA target in each condition was then calculated and normalized to control conditions as outlined (Table 3).

**RNA-seq analysis**

Single-End reads 35-76bp in length were quality checked using fastqc (Andrews, 2010). Reads were mapped to hg38 using the Subread aligner (Liao et al., 2013). The aligned reads were summarized at the gene-level using featureCounts (Liao et al., 2014). Differential gene expression analysis, GO and KEGG pathway enrichment analysis were done using the R/Bioconductor package edgeR (Robinson et al., 2010) and limma (Ritchie et al., 2015).

The average counts-per-million (CPM) values for the EMT gene signatures (cell lines) from (Tan et al., 2014) were computed across the three biological replicates for mCherry$^{NLS}$ (Ctrl), Snail$^{6SA}$ and Snail$^{6SA}$ (extruded) phenotypes (Foroutan et al., 2018). The ternary plot was produced using the R package *vcd*. Specific gene sets for enrichment and visualization were selected based on KEGG pathway membership or through annotation with relevant Gene Ontology terms.

**Antibodies**

The primary antibodies used were as follows: rabbit polyclonal antibody against α-catenin (Invitrogen: #71-1200; IF dilution 1:50); mouse monoclonal antibody against Anillin (Santa Cruz: #sc-271814; IF dilution 1:50 WB dilution 1:500); rabbit polyclonal antibody against ARHGEF28/p190GEF (Life Technologies: #PA558032; WB dilution 1:1000); rabbit polyclonal antibody against ARHGEF31/Ect2 (Merck: #07-1364; IF dilution 1:100 WB dilution 1:1000); mouse monoclonal antibody against ARHGAP5/p190BGAP (BD Bioscience: #611612; IF dilution 1:50 WB dilution 1:1000); rabbit polyclonal antibody against ARHGAP6 (Thermo Fisher Scientific: #PA564170; WB dilution 1:1000); mouse monoclonal antibody against E-cadherin (a gift from P.Wheelock, University of Nebraska, USA; with permission of M.Takeichi; IF dilution 1:50 WB dilution 1:1000); mouse monoclonal antibody against GFP (Rhoche: #11814460001; IF dilution 1:250 WB dilution 1:2000); rabbit polyclonal antibody against GAPDH (Trevigen: #2275; WB dilution 1:10,000); rabbit polyclonal antibody against mCherry (Biovision: #5993; IF dilution 1:250 WB dilution 1:2000); rat monoclonal antibody against mCherry (Thermo Fisher Scientific: #M11217; IF dilution 1:250 WB dilution 1:2000); mouse monoclonal antibody against N-cadherin (Cell Signalling: #14215; WB dilution 1:1000); rabbit polyclonal antibody against Myosin IIA (Sigma: #M8064; IF dilution 1:100 WB dilution 1:1000); rabbit polyclonal antibody against Myosin IIB (Covance: #PRB-445P; IF dilution 1:100 WB dilution 1:1000); rabbit polyclonal antibody against Par3 (Millipore: #07-330; WB dilution 1:1000); rabbit polyclonal antibody against phospho Myosin Light Chain 2 (Ser19) (Cell Signalling: #3675; WB dilution 1:1000); mouse monoclonal antibody against RhoA (Santa Cruz: #SC418; IF dilution 1:50 WB dilution 1:500); rabbit monoclonal antibody against Snail (Cell Signalling: #3879; IF dilution 1:100 WB dilution 1:1000); rabbit monoclonal antibody against Vimentin (Cell Signalling: #5741; WB dilution 1:1000); rabbit polyclonal antibody against ZO-1 (Invitrogen: #61-7300; IF dilution 1:100 WB dilution 1:1000).

Secondary antibodies used for immunoblotting were anti-rabbit (Bio-Rad: #1706515; dilution 1:4000) or anti-mouse (Bio-Rad: #170516; dilution 1:4000) conjugated with horseradish peroxidase (HRP). Secondary antibodies for immunofluorescence were as follows: anti-rabbit AlexaFluor488 (Invitrogen: #A11034; dilution 1:400), anti-mouse AlexaFluor488 (Invitrogen: #A11029; dilution 1:400), anti-rabbit AlexaFluor546 (Invitrogen: #A11035; dilution 1:400), anti-rat AlexaFluor546 (Invitrogen: #A11081; dilution 1:400), anti-rabbit AlexaFluor647 (Invitrogen: #A21245; dilution 1:400) and anti-mouse AlexaFluor647 (Invitrogen: #A21236; dilution 1:400). Phalloidin conjugated with AlexaFluor647 were used to stain for F-actin (Thermo Fisher Scientific: #A22287; dilution 1:400).

**Immunoblotting**

Cells cultured in 6 well plates were placed on ice, washed with ice-cold PBS once, and then lysed with either RIPA lysis buffer or SDS sample lysis buffer. Cells lysates were scraped down, resolved on 10% or 12% SDS-PAGE gels and transferred onto a nitrocellulose membrane. Nitrocellulose membranes were then stained with Ponceau S to visually check equal loading of proteins before blocking with 5% milk diluted in Tris buffered saline with 0.2% tween-20 for 1 hour at room temperature. Specific primary antibodies were then diluted in blocking buffer at the specified dilution as described earlier and added to pre-blocked membranes overnight at 4°C. Membranes were washed with TBST for 3 times, 20 minutes each, at room temperature. Specific secondary antibodies tagged with HRP were diluted in blocking buffer at 1:4000 dilution and incubated with the membrane for 1 hour at room temperature. The membrane was then washed for 3 times, 20 minutes each, at room temperature followed by detection of specific signals by Enhanced Chemiluminescence (ECL) in room temperature. Luminescence was detected in dark and captured on highly sensitive X-films.

**Quantification of Junctional Fluorescence Intensity**

Cells cultured to 95% confluency were fixed with either 4% paraformaldehyde (PFA) in cytoskeletal stabilization buffer for 20 minutes at 37°C, 100% methanol for 5 minutes on ice or 10% TCA/PBS for 20 minutes on ice. For PFA fixation, samples were permeabilised with 0.25% Triton-X-100 for 5 minutes at room temperature. For TCA fixation, samples were washed three times with 30mM Glycine/PBS solution, followed by permeabilisation with 0.25% Triton-X-100. Fixed samples were blocked with 5% milk or 3% BSA prepared in PBS for 1 hour at room temperature, followed by incubation of diluted specific primary antibodies overnight at 4°C in a humid chamber. Samples were then washed with PBS, incubated with diluted specific secondary antibodies for 1 hour in room temperature, and mounted on glass slides with Prolong Gold containing DAPI (Cell Signalling; #8961).

Confocal images were then acquired on Zeiss LSM 710 (63x, 1.4NA Plan Apo objective) using the provided ZEN software. Multiple Z stacks (0.4µM increment) spanning the height of the sample (cells) were acquired for each region that was imaged.

Quantification of fluorescence intensity at junctions was performed using the line scan function in Image J as previously described (Michael et al., 2016). In brief, a line of 20 pixels in width was drawn orthogonal to the cell junction and the pixel intensity was obtained using the Plot Profile function. The peak value for each junction was obtained and corrected by subtracting background fluorescence intensity from either end of the measured line. The resulting value is representative of the fluorescence intensity at the measured junction. Junctional fluorescence intensity from a minimum of 20 different junctions was compiled and analyzed as one independent experiment per condition. The mean peak fluorescence value was then obtained and calculated from three independent experiments and compared between groups.

To quantify junctional active RhoA, the fluorescence intensity of junctional and cytoplasmic GFP-AHPH in $mCherry^{NLS}$ or $Snail^{6SA}$ expressing cells were measured as described above. GFP-AHPH Junctional fluorescence intensity was then corrected for differences in expression levels with the cytoplasmic GFP-AHPH levels for each cell. The average of corrected GFP-AHPH junctional fluorescence values were then compared between groups.

**Quantification of invadopodia punctas**

Homogenous populations of MCF-7 cells expressing either $mCherry^{NLS}$ or $Snail^{6SA}$ were cultured to 95% confluency and fixed with 4% paraformaldehyde as described earlier. Phalloidin staining was used to identify F-actin in the invadopodial structures at the basal region of the cell, beneath the nucleus. Confocal images were then acquired on Zeiss LSM 710 (63x, 1.4NA Plan Apo objective) using the provided ZEN software. Multiple Z stacks (0.4μM increment) spanning the height of the sample (cells) were acquired for each region that was imaged. Invadopodia-like structures were identified as F-actin puncta accumulation beneath the nucleus. F-actin punctas in $Snail^{6SA}$ cells were counted and compared to those of control $mCherry^{NLS}$ cells.

**Quantification of paxillin at focal adhesions**

MCF-7 cells expressing either $mCherry^{NLS}$ or $Snail^{6SA}$ were cultured at low density on glass cover slips to achieve isolated single cells for 5 hours. Cells were then fixed and stained with paxillin for analysis by immunofluorescence. Paxillin punctas around the periphery of $mCherry^{NLS}$ or $Snail^{6SA}$ expressing cells (white box in Figure S6E) were isolated using freehand selections on ImageJ and measured for average fluorescence intensity in the isolated area.

**α-18 and α-catenin ratio analysis**

We utilised the α-18 antibody, a generous gift from the Nagafuchi lab, as a proxy of changes in junctional tension generated by contractile forces at the adherens junction that specifically binds to the ‘exposed’ central domain of the α-catenin protein at cell-cell contacts (Yonemura et al., 2010). Cells were cultured to 95% confluency, fixed with methanol and co-stained with α-18 and α-catenin antibodies at the specified concentration as described earlier. Junctional fluorescence intensity was imaged on confocal microscopy. Junctional fluorescence intensity of α-18 was calculated as a ratio to α-catenin with ratio intensity of $Snail^{6SA}$ cells normalised against those of control $mCherry^{NLS}$ cells.

**Rho-FRET imaging and quantification**

The pTriEx-RhoA Biosensor WT construct was obtained from addgene, deposited by the laboratory of Klaus Hahn (Pertz et al., 2006), and used for RhoA-FRET experiments. pTriEx-RhoA was co-transfected with either $mCherry^{NLS}$ or $Snail^{6SA}$ in MCF-7 cells cultured on glass-bottom dishes. Fresh media containing doxycycline was then replaced the next day for 48 hours to induce expression of $mCherry^{NLS}$ or $Snail^{6SA}$. Live cells were then imaged on Zeiss LSM 710 microscope (63x, 1.40NA Plan Apo objective). Donor and FRET fluorescence channels were excited using the 458nm laser and the emission of donor was collected at a 470-490nm range, while the emission of acceptor molecule during FRET was obtained at a range of 530-590nm as previously described (Liang et al., 2017; Priya et al., 2015).

The images were processed in imageJ to split the fluorescence channels, generate a image stack of various junctions and create a mask of the apical adherens junction for each image to calculate the pixel intensity in various for various channels (Donor, Acceptor and FRET). The Donor and FRET emission at pixels localized at apical cell-cell junctions were used to calculate the emission ratio of FRET/Donor using a custom-made MATLAB script described earlier (Priya et al., 2015; Ratheesh et al., 2012).

**Etoposide-Induced Apoptotic Extrusion Assay**

95% confluent cells were treated with 250µM of etoposide for 5 hrs and analysed by immunofluorescence as described previously (Michael et al., 2016). Cleaved caspase-3 and DAPI were used to identify apoptotic Caco-2 cells while Annexin-V and DAPI were used to identify apoptotic MCF-7 cells. Extruded cells were identified as Annexin-V or cleaved caspase-3 positive spherical cells with their nuclei out of the apical plane of the monolayer. E-cadherin was used to identify the monolayer.

**Extrusion Assay**

To assess apical extrusion, cells stably expressing $mCherry^{NLS}$ or $Snail^{6SA}$ were counted and mixed with wild type cells in a 1:100 ratio and plated at approximately 80% confluency to obtain a mosaic population of a single transgenic cell surrounded by wild type cell. To generate islands/clusters of transgenic cells, $mCherry^{NLS}$ or $Snail^{6SA}$ expressing cells were mixed with wild type cells in a 1:50 ratio and plated at 80% confluency. Transgene expression was then induced by treating with 3 µg/mL of doxycycline for 48 hrs, and the confluent monolayer was fixed for analysis by immunofluorescence. mCherry and DAPI were used to identify extruded cells with their nuclei out of the focus apical plane of the monolayer. E-cadherin was used to identify the apical plane of the monolayer.

For live imaging of apical extrusion, MCF-7 cells in which the endogenous E-cadherin tagged with GFP, using the CRISPR/Cas9 genome editing system, were cultured on glass bottom dishes and transfected with either $mCherry^{NLS}$ or $Snail^{6SA}$ with Lipofectamine 2000 and imaged after 48 hours. Time-lapse live cell imaging of confluent monolayer was performed on the Nikon Ti-E deconvolution microscope (40x, 0.5NA Plan Apo objective), equipped with a 37°C, 5% $CO_2$ chamber, using the NIS-Elements AR software (Nikon). Images were acquired at 30 mins interval for 24 hours.

To assess basal extrusion, cells stably expressing $mCherry^{NLS}$ or $Snail^{6SA}$ were treated with 3 µg/mL of doxycycline for 48 hrs and cultured to approximately 90-100% confluency. E-cadherin was used to identify the apical and baso-lateral regions of the monolayer. mCherry and DAPI immunofluorescence were used to identify basally extruded cells with their nuclei below the basal plane of the monolayer.

For live imaging of basal extrusion, MCF-7 cells expressing endogenous E-cadherin tagged with GFP were infected with either mCherry$^{NLS}$ or Snail$^{6SA}$ viral particles to generate stable cell lines. Time-lapse live cell imaging of cells stably expressing either E-cad-GFP- mCherry$^{NLS}$ or E-cad-GFP-Snail$^{6SA}$ were acquired as described above for live imaging of apical extrusion.

**GTP-RhoA Pull-Down Assay**

Pull-down of GTP–RhoA was performed with the RhoA Pulldown Activation Assay Kit (#BK036, Cytoskeleton) as described previously (Priya et al., 2015). mCherry$^{NLS}$ or Snail$^{6SA}$ expressing cells were treated with 3 µg/mL of doxycycline for 48 hrs and lysed at 95% confluency. The cells were then washed with ice-cold PBS, lysed and processed for the pulldown assay according to the manufacturer's instructions.

**Measurement of Junctional Tension Using Two-Photon Laser Ablation**

The use of two-photon laser to assess junctional tension has been described previously (Leerberg et al., 2014; Liang et al., 2017; Michael et al., 2016; Wu et al., 2014). Adherens junctions of MCF-7 cells were labelled by tagging endogenous E-cadherin with GFP using the CRISPR/Cas9 genome editing system and were used to identify the apical cell-cell contacts within a monolayer. mCherry$^{NLS}$ or Snail$^{6SA}$ were expressed in these genomes edited cells, either stably or transiently for 48hrs with in a confluent monolayer. For the measurement of junctional tension in MRLC$^{AA}$ and MRLC$^{WT}$ cells, MCF-7 cells were transduced with either MRLC$^{AA}$-GFP or MRLC$^{WT}$-GFP lentiviral particles to generate stable cell lines. Junctional localisation of MRLC$^{AA}$ and MRLC$^{WT}$ is used to identify apical cell-cell contacts. All ablation experiments were performed on a Zeiss LSM-510 META confocal microscope (63x, 1.4NA Plan Apo objective) with a 37°C heating stage. Recoil of junctional vertices was recorded by time-lapse imaging at 10 frames with 5s interval between each frame. Analyses of data were performed on ImageJ as previously described (Liang et al., 2017; Michael et al., 2016).

**Table 1 List of primers used in molecular cloning and RT-qPCR**

| Sequence (5' – 3') | Name | Function |
|---|---|---|
| TTACCGGTCGCCACCATGGTGAGCAAG | AgeI-mCherry | Cloning |
| AAATCGATTCAGCGGGGACATCCTGAGCAGC | Snail6SA-ClaI | Cloning |
| AAATCGATTCAGGAGAGCACACACTTGCAGC | HRas-ClaI | Cloning |
| TGTAGTCACGGACTTTCAGGA | Ect2 Fwd | RT-qPCR |
| GTACAATACAACGGGCGACAT | Ect2 Rev | RT-qPCR |
| CCATCATCCTGGTTGGGAAT | RhoA Fwd | RT-qPCR |
| CCATGTACCCAAAAGCGC | RhoA Rev | RT-qPCR |
| GGCGGATTCCATTTGACCTC | P190BGAP Fwd | RT-qPCR |
| ACTATTTCTCGCTGATGCCTACCA | P190BGAP Rev | RT-qPCR |
| CTCTGCTCCTCCTGTTCGAC | GAPDH Fwd | RT-qPCR |
| GCGCCCAATACGACCAAATC | GAPDH Rev | RT-qPCR |
| GGCGATGGCTTCTCTTTCCA | ARHGEF1/p115 Fwd | RT-qPCR |
| ATGATGCTGACGGGAACCAG | ARHGEF1/p115 Rev | RT-qPCR |
| ACAGTTGGTGTGGTAAGTCCA | Anillin Fwd | RT-qPCR |
| CCAGATTCAGCTCGAGGGAC | Anillin Rev | RT-qPCR |
